# Distinct neutrophil effector functions in response to different isolates of *Leishmania aethiopica*

**DOI:** 10.1101/2024.06.27.601019

**Authors:** E. Adem, E. Cruz Cervera, E. Yizengaw, Y. Takele, S. Shorter, J.A. Cotton, G. Getti, P. Kropf

## Abstract

**Background:** In Ethiopia, cutaneous leishmaniasis is mainly caused by *Leishmania* (*L.*) *aethiopica* parasites and presents in three main clinical forms. It is still not clear if the host immune response plays a role in the development of these different presentations. Since neutrophils are likely to be one of the first immune cells present at the site of the sand fly bite, we set up an *in vitro* model of infection of neutrophils with *L. aethiopica* and assessed neutrophil effector functions. We used freshly isolated clinical isolates and one isolate that has been kept in culture for decades.

**Results:** Our results showed by flow cytometry that up to a quarter of neutrophils were associated with *L. aethiopica*; and confocal microscopy demonstrated that all isolates can be internalised. The clinical isolates of *L. aethiopica* associated more efficiently with neutrophils than the long-term cultured *L. aethiopica.* At 18hrs, two distinct populations of neutrophils were identified that associated with *L. aethiopica*, CD15^high^ and CD15^low^ neutrophils.

Our results also showed that all parasites induced apoptosis in *L. aethiopica*-associated neutrophils.

Moreover, our results showed that after 2 hrs, *L. aethiopica*-associated neutrophils upregulated their production of ROS, but to a greater extent with the long-term cultured *L. aethiopica*. After 18 hrs of incubation, CD15^low^parasite^+^ showed an impaired ability to produce ROS as compared to CD15^high^parasite^+^.

**Conclusion:** Using this *in vitro* model, our results show that different *L. aethiopica* parasite isolates, most notably long-term cultured parasites, impacted differently on neutrophil effector functions.

## INTRODUCTION

Cutaneous leishmaniasis (CL) is caused by over 20 different species of *Leishmania* (*L.*) parasites, that are transmitted to their mammalian hosts during the blood meal of infected sand fly vectors. CL, the most common form of leishmaniasis, is endemic in at least 90 different countries. In 2022, over 205,000 cases were reported (1).

In Ethiopia, CL is mainly caused by *L. aethiopica* (2) and presents in three main clinical forms: diffuse cutaneous leishmaniasis (DCL), characterised by numerous non-ulcerating nodules; mucocutaneous leishmaniasis (MCL), where the lesions affect the mucosa of the nose and/or mouth and localised cutaneous leishmaniasis (LCL), characterised by small lesions that progress to ulcers. Whereas LCL usually heals spontaneously, it is not the case for DCL and MCL; both forms are difficult to treat, and relapses are frequent (3).

The mechanisms responsible for the development of these different clinical presentations of CL are not clearly understood. In a recent study, we showed that chemokine and cytokine levels in plasma as well as parasite genetic factors were not associated with different clinical presentations of CL (2). However, only a small number of parasites isolated from DCL and MCL lesions were sequenced, which might explain why we did not identify individual genetic variants significantly associated with disease presentation.

During any infection, neutrophils are key cells of the innate immune response that are quickly recruited following pathogens entry. They possess an array of pathogen recognition molecules, and once engaged, neutrophils can phagocytose and kill microbes by releasing enzymes such as myeloperoxidase and elastase in the phagosome; they can degranulate and release toxic molecules, reactive oxygen species and neutrophil extracellular traps (NETs) that may kill the pathogens in the microenvironment; as well as produce cytokines and chemokines that will promote the recruitment of other immune cells and shape the adaptive immune response (4, 5).

Most of our knowledge of neutrophil effector functions during leishmaniasis is derived from mouse models. It was shown by two-photon microscopy that neutrophils are quickly recruited to the site of sand fly bites (6, 7). Depending on the parasite species, *Leishmania* can be killed by or survive and even multiply in neutrophils (summarised in (8, 9)). The use of neutropenic Genista mice or depletion of neutrophils with monoclonal antibodies at the time of *Leishmania* infection showed that neutrophils can contribute to exacerbation or control of the infection, depending on several factors, such as the route of infection, the genetic background of the mice and the parasite strains (summarised in (8)).

In humans, it has been well documented that neutrophils are quickly recruited to the site of inflammation (12, 13). Since the sand fly bite results in the formation of a pool of blood, neutrophils will be present at the site of infection and further attracted in numbers; they are therefore likely to be one of the first innate immune cells to interact with *Leishmania* parasites.

The interactions between neutrophils and live *L. aethiopica* parasites have not been characterised, and it is not possible to study these interactions by using a mouse model since injection of *L. aethiopica* in mice does not cause symptoms (10, 11).

Therefore, the availability of an *in vitro* cellular model of *L. aethiopica* infection might be useful in identifying differences in neutrophil effector functions in response to the parasites causing different clinical forms of CL.

Here we set up an *in vitro* model of infection of neutrophils with *L. aethiopica* to measure some of the main effector functions of neutrophils and compare these responses between infections with clinical and laboratory isolates of *L. aethiopica*.

## METHODS

### Sample collection

Three ml of blood were collected in heparin tubes from healthy non-endemic controls and was processed immediately after collection: following density gradient centrifugation on Histopaque-1077 (Sigma), neutrophils were isolated from the erythrocyte fraction by dextran sulfate sedimentation as described in (14), resuspended in RPMI containing 5% heat-inactivated fetal bovine serum (FBS)(Sigma)(complete RPMI, cRPMI), 50 IU/mL penicillin and 50 mg/mL streptomycin and used straight away for flow cytometry. Neutrophil purity was >95% and their viability, as determined by 7-aminoactinomycin D (AAD), was >99.0% (Figure S1).

### Leishmania parasites

Three *L. aethiopica* clinical isolates (*L. aethiopica* 1, 2 and 3) were isolated from the lesions of three different LCL patients from Ethiopia as described in (2) and frozen. To confirm that these isolates were *L. aethiopica*, we mapped RNA-seq reads from the isolates to the reference genome for *L. aethiopica* (15) obtained from TriTrypdb using STAR v2.7.0 (16), and then used samtools v1.17 (17) to call the consensus sequence of these mapped reads over the HSP70 locus (LAEL147_000511500), which has been widely used to speciate *Leishmania* isolates. We compared these reconstructed sequences with previously obtained sequences from a range of *Leishmania* species (principally from (18)) using mafft v7.45 (19) to align the sequences, trimAl v2.0 (20) (with flag-strictplus) and then building a phylogeny using raxmlHPC v8.2.12 (21) (a single tree search under a GTR+gamma model of nucleotide substitution). This analysis confirmed that the isolates used here have sequences identical to *L. aethiopica* (accession FN395019) and differing by a single SNP from two other *L. aethiopica* isolates (FN395020 and FN395021). Other species formed separate groups on a phylogeny based on these sequences.

Another isolate of *L. aethiopica* (MHOM/ET/72/L100)(22, 23) was also used. Even though it is not known precisely how long and how many times it had been kept frozen and in culture, it is known that it was isolated over 40 years ago (10); this long-term cultured *L. aethiopica* isolate was therefore identified as *L. aethiopica* “laboratory” (*L. aethiopica* lab). A large stock of frozen stationary phase *L. aethiopica* 1, 2 and 3 and lab were prepared for further analysis. Once thawed, the parasites were used for a maximum of 3 weeks.

The following culture medium was used to keep the parasites in culture: M199 medium with 25mM hepes, 0.2μM folic acid, 1mM hemin, 1mM adenine, 800μM Biopterin, 50 IU/mL penicillin, 50 mg/mL streptomycin and 10% FBS (Sigma, USA) and the parasites were incubated at 26°C.

Metacyclic promastigotes were isolated via agglutination with peanut agglutinin (PNA) as described previously (24) and were stained using CellTrace™ Far Red dye (FR) (Invitrogen, UK), using a 1 µM FR solution. Following a 20-minute incubation at room temperature, the suspension was diluted 1 in 10 in cRPMI medium and incubated for further 5 minutes at room temperature to quench any free dye remaining in the solution. At the end of the incubation, parasites were washed and resuspended in cRPMI medium and used for further experiments.

### Neutrophil effector functions

Human neutrophils (1×10^5^ cells/ml) were co-cultured with 1×10^6^ cells/ml FR labelled *L. aethiopica* isolates for 2 hours, at 37°C, 5% CO_2_, in cRPMI. Cells were then washed twice with PBS, and cells to be used for the 2 hrs incubation were labelled with antibodies as described below for flow cytometry analyses. Cells for the 18 hrs incubation were resuspended in cRPMI and incubated for a further 16 hours at 37°C, 5% CO_2_, washed twice with PBS and processed as indicated above for the 2 hrs incubation.

#### Association of parasites with neutrophils

The % of association between neutrophils (stained with anti-human CD15^eFluor^ ^450^ (clone MMA) (eBioscience) and FR labelled *L. aethiopica* was assessed by flow cytometry.

#### Confocal microscopy

After 2 and 18 hrs of incubation, cells were transferred to poly-L-lysine (0.01% solution, Sigma-Aldrich) coated coverslips and were incubated at room temperature. After 30 minutes, the cells were washed twice with PBS and fixed with 2% (w/v) paraformaldehyde (Sigma-Aldrich, UK) for 20 min. Cells were then washed three times with PBS and incubated with CD15 (C3D-1) mouse anti-human monoclonal antibody (Thermo Fisher Scientific) overnight at 4°C, followed by anti-IgG (H+L) highly Cross-Adsorbed secondary antibody (Alexa Fluor 555) (Thermo Fisher Scientific). After one hr of incubation in the dark, the coverslips were washed three times with PBS and placed onto a slide containing 50 µl of mounting media (VECTASHIELD mounting media, Vector Laboratories). Slides were visualised under a ZEISS LSM 880 Confocal Laser Scanning Microscope under 60X magnification. 1.00 Airy units (1AU) pinhole size was used. Image acquisition was done using Zen black software and the 3D z-stack orthogonal images were analysed by Zen 3.3 (blue edition) software.

#### Apoptosis

The PE-Annexin V/7-amino-actinomycin D (7-AAD) apoptosis detection kit (BioLegend, UK) was used to detect apoptosis according to the manufacturer’s protocol.

#### ROS detection assay

ROS-ID^TM^ Total ROS detection kit (Enzo Life Sciences, USA) was used to evaluate the production of ROS by neutrophils according to the manufacturer’s protocol.

Flow cytometry acquisition was performed using an LSRII (BD Biosciences) and data were analysed using Summit v4.3 software.

### Statistical analysis

Data were evaluated for statistical differences as specified in the legend of each figure. The following tests were used: Mann-Whitney and Kruskal-Wallis. Results are expressed as mean ± SD. Differences were considered statistically significant at *p*<0.05. *=p<0.05, **=p<0.01, ***=p<0.001 and ****=p<0.0001.

## RESULTS

### Association of *L. aethiopica* parasites with neutrophils

We first assessed by flow cytometry whether the three clinical and the laboratory isolates of *L. aethiopica* can associate with neutrophils after 2 hrs of co-incubation. Results presented in Figures 1A and B show that *L. aethiopica* can associate with neutrophils, and that the percentages of the *L. aethiopica* lab associated with neutrophils were significantly lower as compared to the three clinical isolates (Figure 1B and Table 1). There were no significant differences between the three clinical isolates (Figure 1B and Table 1).

**Figure 1:**
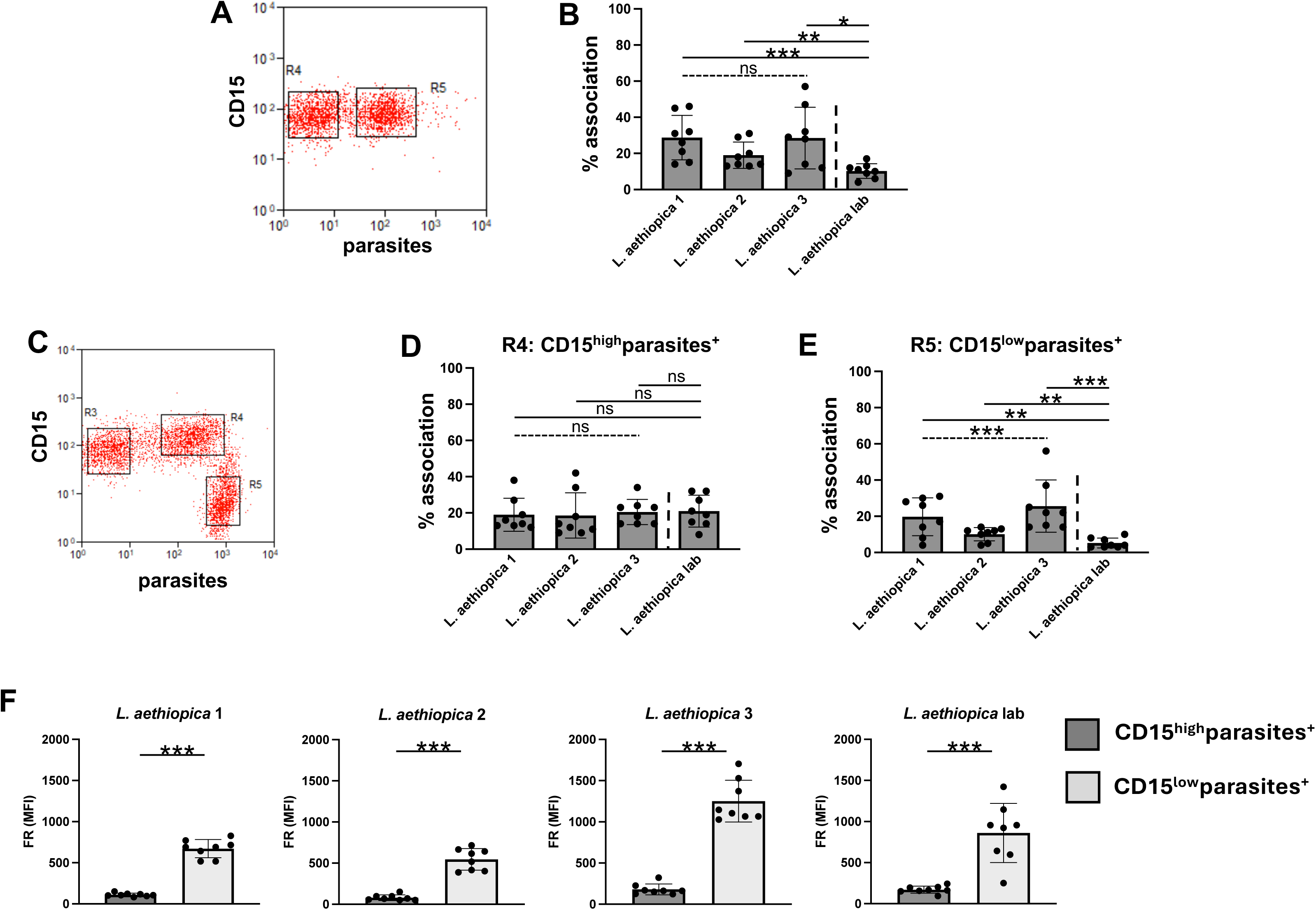
Association of neutrophils with different isolates of *L. aethiopica*. 1×10^5^ cells/ml neutrophils were co-cultured with 1×10^6^ cells/ml FR labelled *L. aethiopica* isolates for 2 hrs (**A** and **B**) and 18 hrs (**C**, **D**, **E** and **F**). The percentages of neutrophils associated with *L. aethiopica* were determined by flow cytometry. **A.** Dot plot showing neutrophils unassociated (gate R4) and associated (gate R5) with *L. aethiopica*. **B**. % of neutrophils associated with the four different isolates of *L. aethiopica*. **C.** Dot plot showing the three different population of neutrophils: CD15^intermediate^ ^(int)^parasite^-^ (gate R3), CD15^high^parasite^+^ (gate R4) and CD15^low^parasite^+^ (gate R5). **D.** % of CD15^high^parasite^+^ associated with the four different isolates of *L. aethiopica.* **E.** % of CD15^low^parasite^+^ associated with the four different isolates of *L. aethiopica.* **F.** Comparison in FR MFI between CD15^high^parasite^+^ and CD15^low^parasite^+^ for each *L. aethiopica*. Data are presented as scatter plot with bar (mean with standard deviation), with each dot representing the value for one experiment. Statistical differences were determined using Kruskal-Wallis (dotted line) and Mann-Whitney (solid line) tests.

**Table 1:** Association of neutrophils with *L. aethiopica* following 2 hrs of co-incubation.

| Clinical isolates | % association | *p value |  |
| --- | --- | --- | --- |
| *L. aethiopica* 1 | 28.8±12.6 | 0.2425 |  |
| *L. aethiopica* 2 | 19.0±7.3 |  |  |
| *L. aethiopica* 3 | 28.5±17.0 |  |  |
|  |  | *L. aethiopica* lab<br>% association | #p value |
| *L. aethiopica* 1 | 28.8±12.6 | 10.3±6.9 | 0.0006 |
| *L. aethiopica* 2 | 19.0±7.3 |  | 0.0023 |
| *L. aethiopica* 3 | 28.5±17.0 |  | 0.0134 |

1x10<sup>5</sup> cells/ml neutrophils were co-cultured with 1x10<sup>6</sup> cells/ml FR labelled *L. aethiopica* isolates for 2 hours. The percentages of neutrophils associated with *L. aethiopica* were determined by flow cytometry. Statistical differences were determined using Kruskal-Wallis (\*) and Mann-Whitney (#) tests.

Since it has been shown that infection of neutrophils with *Leishmania* parasites delays apoptosis for up to 24 hrs (25), we assessed whether their ability to associate with parasites was maintained. As shown in Figure 1C, there were two distinct populations of neutrophils associated with the parasites: CD15^high^ (gate R4) and CD15^low^ (gate R5). This was the case for all 4 isolates of *L. aethiopica* (data not shown). There were no significant differences between different isolates in the percentages of parasites associated with neutrophils in the CD15^high^parasites+ (Figure 1D and Table 2). However, the percentages of CD15^low^ cells associated with *L. aethiopica* 2 were significantly lower as compared to *L. aethiopica* 3 (Figure 1E and Table 3). Furthermore, the percentages of the three clinical isolates associated with CD15^low^ cells were all significantly higher as compared to *L. aethiopica* lab (Figure 1E and Table 3). Since there were differences in FR intensities between the four parasite isolates (Figure S2), it was not possible to compare the intensity of association between the neutrophils and the different isolates. However, it was possible to compare between the CD15^high^ and the CD15^low^ populations for each parasite isolate. As shown in Figure 1F, the MFI of the FR labelled parasites were always significantly higher in the CD15^low^ population, suggesting that more parasites were associated with the CD15^low^ population.

**Table 2:** Comparison of the association of CD15^high^ neutrophils with the different *L. aethiopica* isolates after 18 hrs.

| CD15<sup>high</sup>parasites<sup>+</sup> |  |  |  |
| --- | --- | --- | --- |
| Clinical isolates | % association | *p value |  |
| *L. aethiopica* 1 | 19.0±9.1 | 0.3728 |  |
| *L. aethiopica* 2 | 18.6±12.5 |  |  |
| *L. aethiopica* 3 | 20.5±7.0 |  |  |
|  |  | *L. aethiopica* lab<br>% association | #p value |
| *L. aethiopica* 1 | 19.0±9.1 | 21.0±8.8 | 0.4525 |
| *L. aethiopica* 2 | 18.6±12.5 |  | 0.4559 |
| *L. aethiopica* 3 | 20.5±7.0 |  | 0.8973 |

1x10<sup>5</sup> cells/ml neutrophils were co-cultured with 1x10<sup>6</sup> cells/ml FR labelled *L. aethiopica* isolates for 18 hours. The percentages of CD15<sup>high</sup> neutrophils associated with *L. aethiopica* were determined by flow cytometry. Statistical differences were determined using Kruskal-Wallis (\*) and Mann-Whitney (#) tests.

**Table 3:**
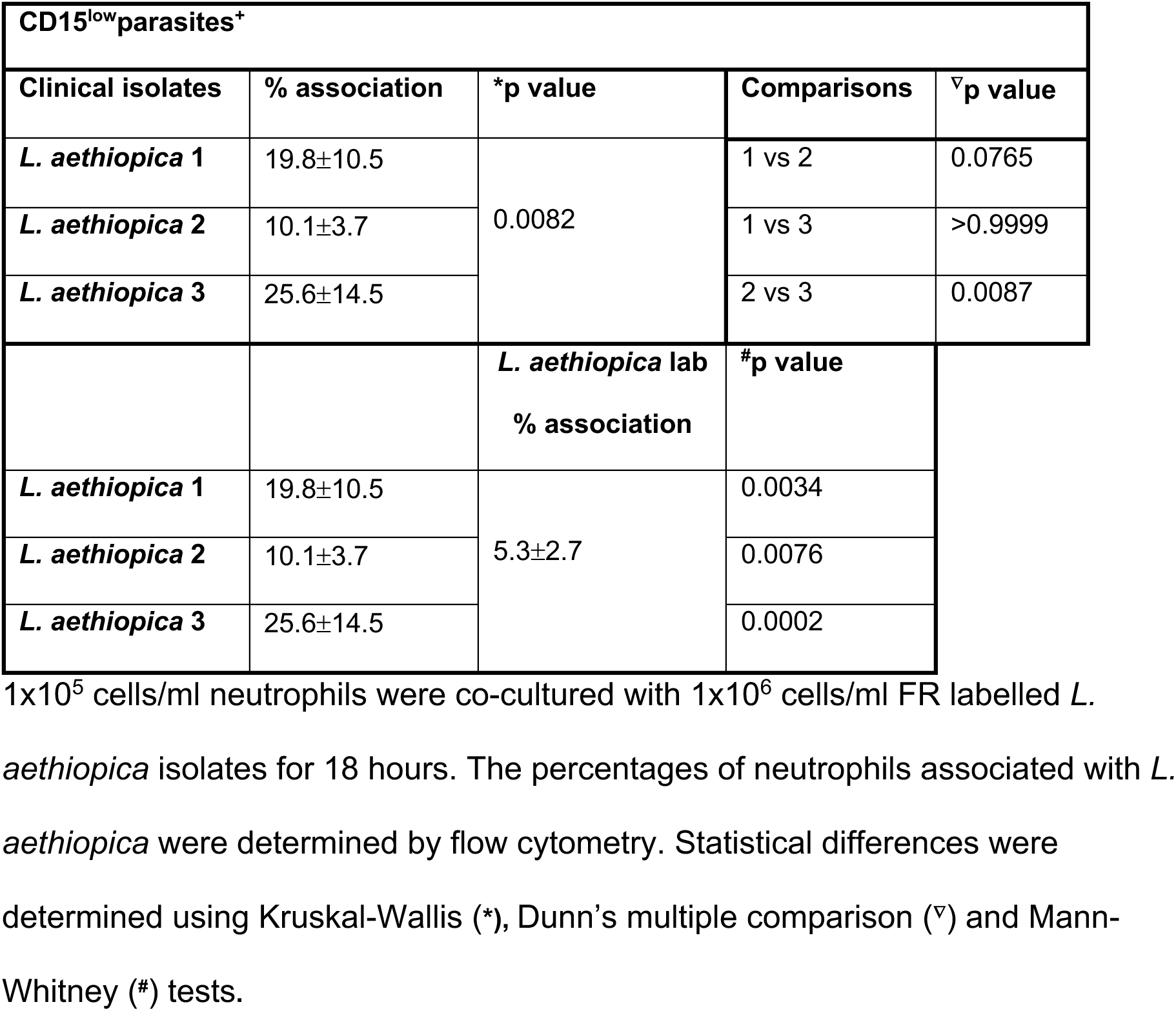
Comparison of the association of CD15^low^ neutrophils with the different *L. aethiopica* isolates after 18 hrs.

| CD15<sup>low</sup>parasites<sup>+</sup> |  |  |  |  |
| --- | --- | --- | --- | --- |
| Clinical isolates | % association | *p value | Comparisons | ▽p value |
| *L. aethiopica* 1 | 19.8±10.5 | 0.0082 | 1 vs 2 | 0.0765 |
| *L. aethiopica* 2 | 10.1±3.7 |  | 1 vs 3 | >0.9999 |
| *L. aethiopica* 3 | 25.6±14.5 |  | 2 vs 3 | 0.0087 |
|  |  | *L. aethiopica* lab<br>% association | #p value |  |
| *L. aethiopica* 1 | 19.8±10.5 | 5.3±2.7 | 0.0034 |  |
| *L. aethiopica* 2 | 10.1±3.7 |  | 0.0076 |  |
| *L. aethiopica* 3 | 25.6±14.5 |  | 0.0002 |  |

1x10<sup>5</sup> cells/ml neutrophils were co-cultured with 1x10<sup>6</sup> cells/ml FR labelled *L. aethiopica* isolates for 18 hours. The percentages of neutrophils associated with *L. aethiopica* were determined by flow cytometry. Statistical differences were determined using Kruskal-Wallis (\*), Dunn's multiple comparison (▽) and Mann-Whitney (#) tests.

### Internalisation of *L. aethiopica* by neutrophils

Next, confocal microscopy was used to demonstrate that *L. aethiopica* is internalised by neutrophils and does not only associate as shown by flow cytometry. Results presented in Figure 2 (*L. aethiopica* lab) and Figures S3 (*L. aethiopica* 1, 2 and 3) show that *L. aethiopica* parasites were internalised within neutrophils following 2 and 18 hrs of incubation. The top panel and the right panel, both delineated by a grey line, show the horizontal and vertical view of the z-stack. Both show that the internalised parasites are surrounded by CD15^+^ neutrophil membrane.

**Figure 2:**
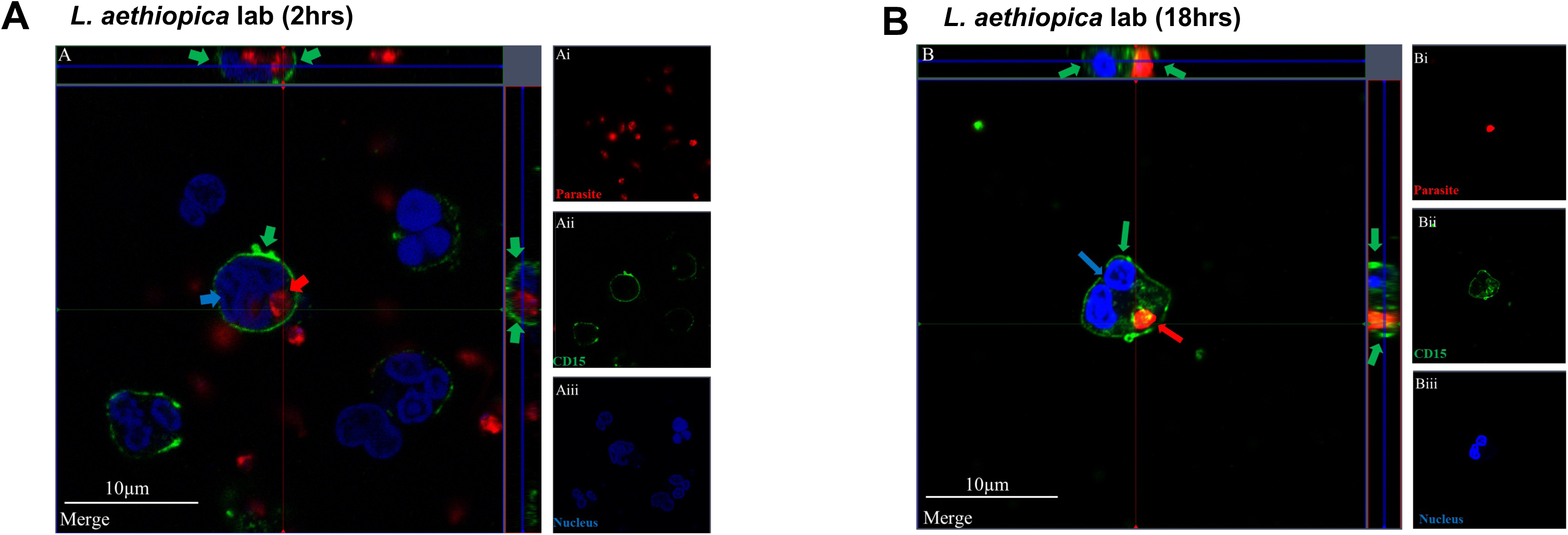
Internalisation of *L. aethiopica* by neutrophils. 1×10^5^ cells/ml neutrophils were co-cultured with 1×10^6^ cells/ml FR labelled *L. aethiopica* lab for 2 hrs (A) and 18 hrs (B) and cells were labelled as described in Materials and Methods. The red arrows point to the parasite (FR), the green arrows to the CD15 (Alexa Fluor 555) and the blue arrow to the nucleus (DAPI). These are representative images of at least three independent experiments.

### Neutrophil apoptosis

To determine how co-culture of neutrophils with *L. aethiopica* impacts on their ability to undergo apoptosis, we measured the percentages of apoptotic cells (as defined by Annexin V^+^ 7-AAD^-^ (early apoptosis) or Annexin V^+^ 7-AAD^+^ (late apoptosis) neutrophils. The gating strategy is shown in Figure S4. As shown in Figures S4 D and E, the majority of gated apoptotic neutrophils were in the early stage of apoptosis. Following 2 hrs of incubation, the percentages of apoptotic cells were systematically increased in neutrophils associated or not with all *L. aethiopica* isolates as compared to neutrophils alone (Table S1). Of note, the % of apoptotic neutrophils were significantly higher in neutrophils associated with the three clinical isolates as compared to unassociated neutrophils; this was however not the case with *L. aethiopica* lab (Table S2).

To be able to compare the % of neutrophils undergoing apoptosis following incubation with the different parasite isolates, % changes in apoptosis were assessed between baseline (neutrophils incubated without parasites) and those neutrophils co-incubated with *L. aethiopica*. Results show that after 2 hrs of incubation, there was no significant difference between any isolates in % change in apoptosis by neutrophils associated (Figure 3A) or not (data not shown) with *L. aethiopica* between all isolates.

**Figure 3:**
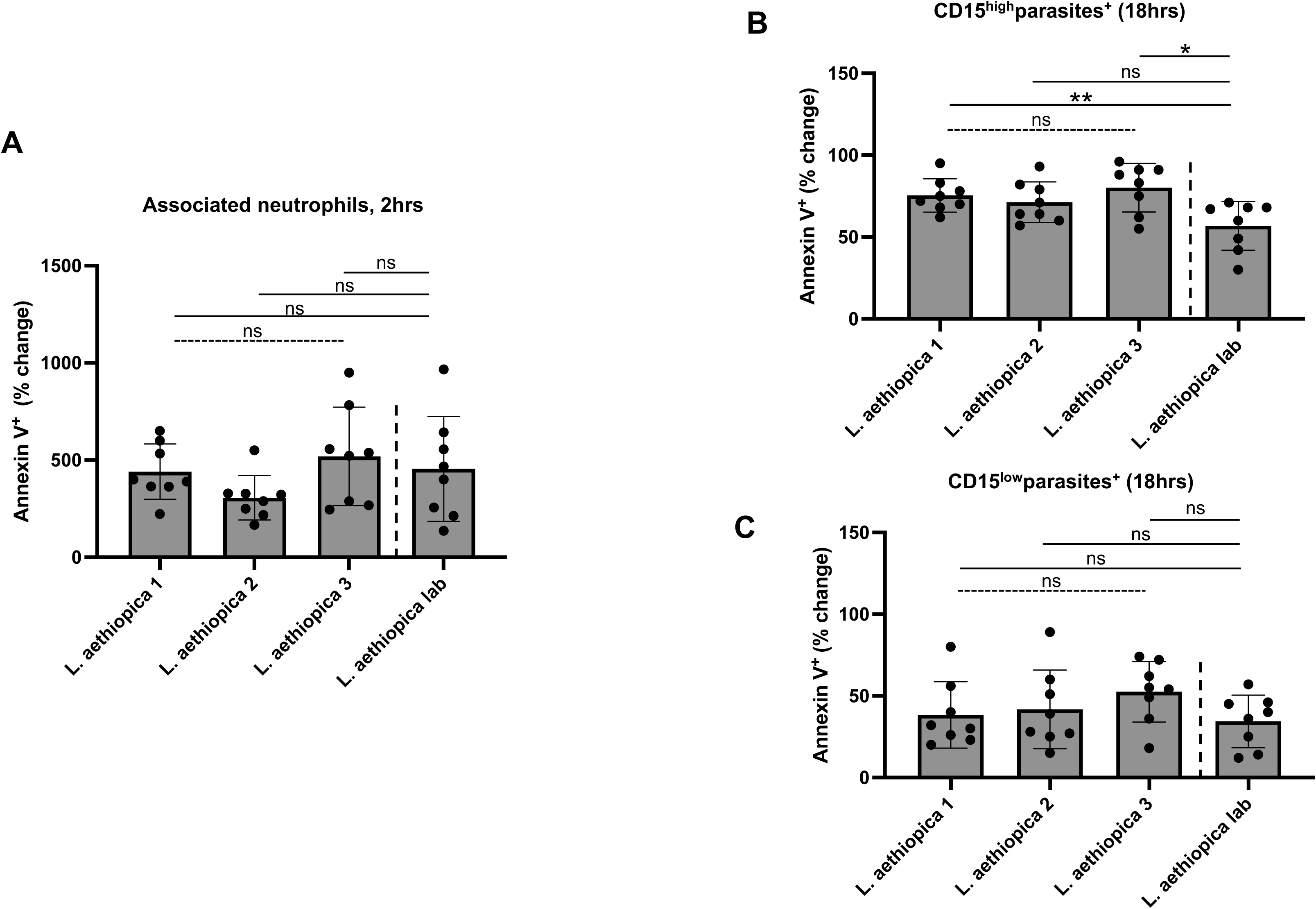
% change in apoptotic neutrophils between the different parasite isolates. 1×10^5^ cells/ml neutrophils were co-cultured with 1×10^6^ cells/ml FR labelled *L. aethiopica* isolates for 2 hrs (**A**) and 18 hrs (**B** and **C**). The % change was measured by deducting the % apoptotic (as defined by Annexin V^+^7-AAD^-^) CD15^high^ neutrophils co-cultured with the parasites from the % of apoptotic neutrophils cultured in the absence of parasites. Data are presented as scatter plot with bar (mean with standard deviation), with each dot representing the value for one experiment. Statistical differences were determined using Kruskal-Wallis (dotted line) and Mann-Whitney (solid line) tests.

After 18 hours of incubation (gating strategy shown in Figure S5), the majority of gated apoptotic neutrophils were in the early stage of apoptosis (Figure S5). Results presented in Table S3 show that for all isolates, the % Annexin V^+^ 7-AAD^-^ neutrophils were similar between all four parasite isolates in CD15^int^parasites^-^ and lower in CD15^high^parasites^+^ and CD15^low^parasites^+^ as compared to baseline. When comparing the % Annexin V^+^ 7-AAD^-^ neutrophils between the three population of neutrophils, it was always highest in the CD15^int^parasites^+^ and lowest in the CD15^low^parasites^+^ (Table S4).

There were no significant differences in % change in the frequency of apoptotic cells between the 4 different isolates in CD15^int^parasite^-^ (data not shown). In the CD15^high^parasites^+^, the % change was significantly higher with *L. aethiopica* 1 and 3, but not 2, as compared to *L. aethiopica* lab (Figure 3B and Table 4). However, there were no significant differences in % change between any parasites isolates in the CD15^low^parasites^+^ (Figure 3C).

**Table 4:** % change in apoptotic CD15^high^ neutrophils between the different parasite isolates after 18 hrs.

|  | % change | *p value |  |
| --- | --- | --- | --- |
| *L. aethiopica* 1 | 75±10 | 0.3950 |  |
| *L. aethiopica* 2 | 71±12 |  |  |
| *L. aethiopica* 3 | 80±15 |  |  |
|  |  | *L. aethiopica* lab<br>% change | #p<br>value |
| *L. aethiopica* 1 | 75±10 | 56.88±15 | 0.0067 |
| *L. aethiopica* 2 | 71±12 |  | 0.1514 |
| *L. aethiopica* 3 | 80±15 |  | 0.0135 |

1x10<sup>5</sup> cells/ml neutrophils were cultured in the presence or the absence of 1x10<sup>6</sup> cells/ml FR labelled *L. aethiopica* isolates for 18 hrs. The % change was measured by deducting the % apoptotic (as defined by Annexin V<sup>+</sup>7AAD<sup>-</sup>) CD15<sup>high</sup> neutrophils co-cultured with the parasites from the % of apoptotic neutrophils cultured in the absence of parasites. Statistical differences were determined using Kruskal-Wallis (\*) and Mann-Whitney (#) tests.

### ROS production by *L. aethiopica*-associated neutrophils

Next, we assessed the ability of neutrophils to upregulate ROS following co-incubation with *L. aethiopica* (gating strategy shown in Figure S6). Following 2 hrs of incubation, ROS production (MFI) was systematically significantly increased in *L. aethiopica*-associated neutrophils, but it was not significantly higher in unassociated neutrophils, except for a borderline difference with *L. aethiopica* 2 (p=0.0433, Table S5). Of note, the levels of ROS production (MFI) were significantly higher in *L. aethiopica*-associated neutrophils than in unassociated ones (Table S6).

To be able to compare the levels of ROS production by neutrophils between the different parasite isolates, % changes in ROS production were assessed between baseline (neutrophils incubated without parasites) and those neutrophils co-incubated with *L. aethiopica*. Results shown in Figure 4A and Table 5 show that after 2 hrs of incubation, there was no significant difference in % change in ROS production by neutrophils associated with *L. aethiopica* between the three clinical isolates. However, it was significantly higher with *L. aethiopica* lab as compared to the clinical isolates (Figure 4A and Table 5). There was no significant difference in ROS production between all four isolates in the *L. aethiopica*-unassociated neutrophils (data not shown).

**Figure 4:**
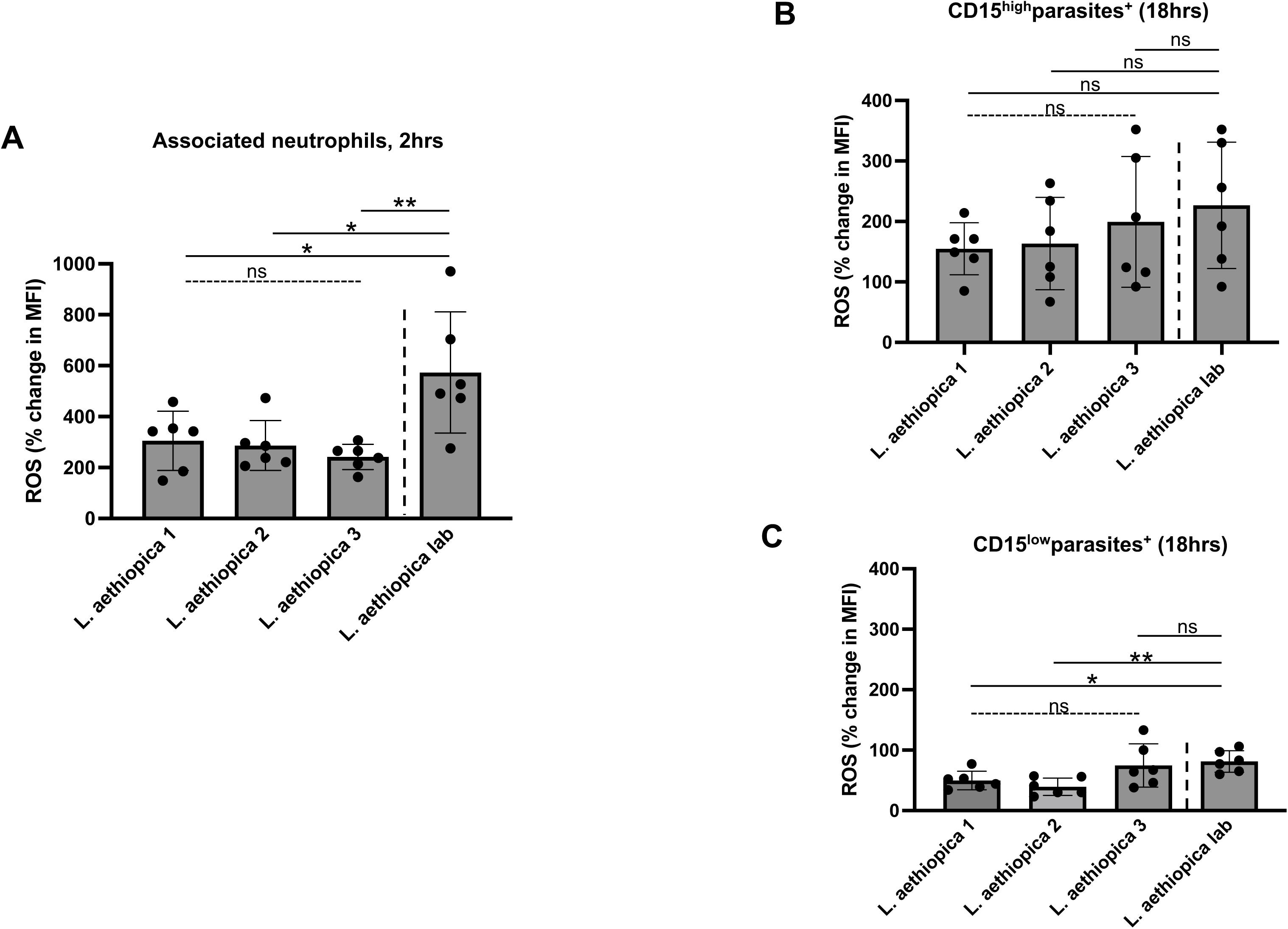
% change in ROS MFI in neutrophils between the different parasite isolates. 1×10^5^ cells/ml neutrophils were co-cultured with 1×10^6^ cells/ml FR labelled *L. aethiopica* isolates for 2 hrs (**A**) and 18 hrs (**B** and **C**). The % change was measured by deducting the ROS MFI in the neutrophils co-cultured with the parasites from the ROS MFI in the neutrophils cultured in the absence of parasites. Data are presented as scatter plot with bar (mean with standard deviation), with each dot representing the value for one experiment. Statistical differences were determined using Kruskal-Wallis (dotted line) and Mann-Whitney (solid line) tests.

**Table 5:** % change in ROS MFI in neutrophils between the different parasite isolates after 2 hrs.

| ROS | % change | *p value |  |
| --- | --- | --- | --- |
| *L. aethiopica* 1 | 305±16 | 0.5885 |  |
| *L. aethiopica* 2 | 287±98 |  |  |
| *L. aethiopica* 3 | 242±50 |  |  |
| ROS |  | *L. aethiopica* lab<br>% change | #p value |
| *L. aethiopica* 1 | 305±16 | 573±238 | 0.0216 |
| *L. aethiopica* 2 | 287±98 |  | 0.0173 |
| *L. aethiopica* 3 | 242±50 |  | 0.0043 |

1x10<sup>5</sup> cells/ml neutrophils were cultured in the presence or the absence of 1x10<sup>6</sup> cells/ml FR labelled *L. aethiopica* isolates for 2 hrs. The % change was measured by deducting the ROS MFI in the neutrophils co-cultured with the parasites from the ROS MFI in the neutrophils cultured in the absence of parasites. Statistical differences were determined using Kruskal-Wallis (\*) and Mann-Whitney (#) tests.

After 18 hours of incubation (gating strategy shown in Figure S7), results presented in Table S7 show that ROS MFI was similar when comparing baseline with CD15^int^parasites^-^ and CD15^high^parasites^+^ neutrophils for all four *L. aethiopica* isolates. However, ROS MFI was significantly lower in CD15^low^parasites^+^ as compared to baseline for all three clinical isolates, but not *L. aethiopica* lab (Table S7). When comparing ROS production between the three populations of neutrophils, the CD15^high^parasites^+^ always produced the highest levels of ROS as compared to CD15^low^parasites^+^ (Table S8). There were no significant differences in % change between the 4 different isolates in CD15^high^parasite^+^ (Figure 4B) and CD15^int^parasite^-^ (data not shown). However, the % change in ROS production in CD15^low^parasite^+^ neutrophils were lower with *L. aethiopica* 1 and 2, but not 3, as compared to *L. aethiopica* lab (Figure 4C and Table 6).

**Table 6:** % change in ROS MFI in CD15^low^ neutrophils between the different parasite isolates after 18 hrs.

| ROS | % change | *p value |  |
| --- | --- | --- | --- |
| *L. aethiopica* 1 | 50±15 | 0.0908 |  |
| *L. aethiopica* 2 | 40±14 |  |  |
| *L. aethiopica* 3 | 75±36 |  |  |
| ROS |  | *L. aethiopica* lab<br>% change | #p value |
| *L. aethiopica* 1 | 50±15 | 81±18 | 0.0108 |
| *L. aethiopica* 2 | 40±14 |  | 0.0022 |
| *L. aethiopica* 3 | 75±36 |  | 0.5887 |

1x10<sup>5</sup> cells/ml neutrophils were cultured in the presence or the absence of 1x10<sup>6</sup> cells/ml FR labelled *L. aethiopica* isolates for 18 hrs. The % change was measured by deducting the ROS MFI in the CD15<sup>low</sup> neutrophils co-cultured with the parasites from the ROS MFI in the CD15<sup>low</sup> neutrophils cultured in the absence of parasites. Statistical differences were determined using Kruskal-Wallis (\*) and Mann-Whitney (#) tests.

## DISCUSSION

Here we set up an *in vitro* model of infection of human neutrophils with *L. aethiopica* and show that parasites can be phagocytosed and modulate apoptosis and ROS production. Our results also show that *L. aethiopica* lab associated less and induced more ROS production as compared to freshly isolated *L. aethiopica*. We also identify some differences in the percentages of associated parasites between the three clinical isolates.

Little is known about the mechanisms influencing the different clinical presentations of cutaneous lesions caused by *L. aethiopica*. It has been previously suggested that differences in parasites are associated with the different clinical manifestations (26–28). A later study suggested that the genetic variability did not correlate with the different manifestations (29). In agreement with these results, in the largest study to date analysing genetic variations between *L. aethiopica* isolated from different lesions, we found no individual genetic variants were significantly associated with disease presentation (2).

Here we used three *L. aethiopica* parasites freshly isolated from patients with localised cutaneous leishmaniasis (LCL); and one isolate that had been kept in culture for decades; the history of the number of *in vitro* passages as well as its origin have been poorly characterised (/documented?) (10). *L. aethiopica* parasites cannot be passaged *in vivo* (10, 11) to maintain virulence (30–32). Therefore, to minimise variations due to the time in culture, once the parasites were growing from the cultured skin scrapings from CL patients, they were immediately frozen and shipped to the UK. Large stocks of parasites were produced and frozen, and once thawed, the parasites were kept in culture for a maximum of 3 weeks, before a new tube was thawed. It has been shown that the metacyclogenesis of parasites is key in determining the virulence of the parasites (33). Therefore, to maximise our control over the stage of the parasites used in these experiments, parasites were grown to a stationary phase and metacyclic parasites were purified using PNA.

Long term *in vitro* culture has been associated with the loss of virulence of *Leishmania* parasites both in phagocytic cells, as shown by less amastigotes per cells (30–32); and in animal models, as shown by a lower parasite burden or smaller lesions (32, 34, 35). Several virulence factors have been identified in *Leishmania* parasites (36). In particular, reduced expression of lipophosphoglycan (LPG) and glycoprotein (GP)63 have been shown to result from long-term *in vitro* culture (37, 38). Both these molecules are important in the phagocytosis of *Leishmania* parasites (39, 40). This might therefore explain why *L. aethiopica* lab did associate significantly less with neutrophils, as compared to the freshly isolated *L. aethiopica* parasites.

Interestingly, following 18 hrs of incubation, in addition to a population of neutrophils that did not associate with *L. aethiopica*, two other populations of neutrophils that were associated with *L. aethiopica* were identified; based on the expression levels of CD15. These results show that following co-incubation of neutrophils with *L. aethiopica*, at least three populations of neutrophils can be identified, that have different abilities to associate with the parasites. The heterogeneity of neutrophils is now well recognised (5, 41, 42). scRNAseq analysis of circulating neutrophils showed three major populations (43). Another study showed a high level of heterogeneity of neutrophils following phenotypic characterisation of peripheral neutrophils in healthy individuals and as compared to patients with different pathological conditions (44). We also found differences in the percentages of associated CD15^low^ neutrophils between the three clinical isolates at 18 hrs; these results suggest that even though all three clinical isolates were obtained from lesions of LCL patients, there still might be differences between these parasites.

Infection of neutrophils by *L. major* and *L. infantum* has been shown to result in delayed apoptosis over time, suggesting that the parasites prolong the survival of neutrophils, thereby allowing the intracellular parasites to survive longer (25, 45). Our results show that after 2 hrs of incubation, there were increased percentages of apoptotic associated and unassociated neutrophils as compared to baseline. This was in contrast to 18hrs, when there were significantly lower percentages of apoptotic associated neutrophils (CD15^high^ and CD15^low^), but not unassociated neutrophils (CD15^int^), as compared to baseline. The study by van Zandbergen *et al*. showed that apoptotic neutrophils can be phagocytosed by macrophages and that the phagocytosed parasites are then able to survive and multiply in these macrophages (46). It is therefore possible that it will also be the case with *L. aethiopica* and that apoptotic neutrophils will be taken up by phagocytic cells, monocytes in the blood and macrophages if they enter tissues. It has been previously shown that as compared to uninfected neutrophils, *L*. *major*-infected neutrophils have an increased ability to take up noninfected apoptotic cells (47); this was associated with the downregulation of ROS production and better survival of parasites in the neutrophils. Thus, the high percentages of unassociated neutrophils identified in our study might contribute to a better survival of the intracellular parasites. Whereas it has been shown that *Leishmania*-infected neutrophils undergo apoptosis (25, 45–47), there is little information about unassociated neutrophils undergoing apoptosis. They might become apoptotic as a result of a transient contact with *Leishmania* parasites, induction of ROS or the production of cytokines such as TNFα (summarised in (48)).

In our study, after 18 hrs, there were fewer apoptotic cells in associated neutrophils as compared to baseline. This is in contradiction with the study by Oualha *et al.*, as they show an increase in the percentages of apoptotic neutrophils following co-incubation with *L. major* and *L. infantum* as compared to baseline. This might be due to differences in the experimental conditions, such as different parasite strains, a ten-fold higher number of neutrophils co-cultured with parasites and differences in the stage of the parasites, as in this study, they did not use PNA to isolate the metacyclic parasites (45).

It has been previously shown that infection of neutrophils with different parasite species such as *L. infantum* (49, 50), *L. braziliensis* (51) and *L. donovani* (52) results in upregulation of ROS. One study showed higher levels of ROS by neutrophils in response to the parasites; however, lysate and not live *L. aethiopica* were used in this study (53). In our study, ROS was upregulated in neutrophils following 2 hrs of co-culture with all parasite isolates. In contrast, after 18 hrs, ROS was similar to baseline levels in CD15^int^parasites^-^ and in CD15^high^parasites^+^ neutrophils, and significantly lower in CD15^low^parasites^+^. The latter subpopulation was also the population that had the highest % of associated parasites, suggesting that the parasites might manipulate the neutrophils to limit the levels of ROS production to survive more efficiently. In support of these data, ROS had already been shown to be reduced following exposure of *L. major*-infected neutrophils to apoptotic cells (47).

Furthermore, a study by Mollinedo *et al.* has shown that *L. major*- and *L. donovani*-containing phagosomes do not fuse with specific and tertiary granules, thereby preventing the production of ROS (54). Al Tuwaijri *et al.* have also shown that different preparations of *L. major* parasites reduced the respiratory burst by neutrophils (55).

Of note, *L. aethiopica* lab induced higher levels of ROS at both 2 and 18 hrs as compared to the clinical isolates. Both LPG (56) and GP63 (57) have been shown to inhibit the oxidative burst. As discussed above, the extensive time in culture might have resulted in lower expression levels of LPG and GP63 on *L. aethiopica* lab and thereby impacted the levels of ROS production.

### Conclusions

In this study we characterised for the first time neutrophil effector functions in response to different isolates of *L. aethiopica*. Our results show that care should be taken when using parasites that have been kept in culture for a long time and highlight that a more standardised way to isolate primary cells and infectious metacyclic parasites should be used.

In the absence of a mouse model, *in vitro* infection of phagocytic cells might identify different functional profiles that might shed light on the different presentation of lesions caused by *L. aethiopica*.

## DECLARATIONS

### Ethical approvals

Ethical approval was obtained from the Faculty Ethic Committee University of Greenwich (FES-FREC-20-01.04.08.CA). Informed written consent was obtained from each participant.

### Consent for publication

Not applicable

### Availability of data and materials

The datasets supporting the conclusions of this article are included within the article (and its additional file(s))

### Authors’ contributions

Conception of the study: EA, GG, PK

Acquisition of the data: EA, ECC, EY, YT, PK

Analysis: EA, ECC, YT, SS, JAC, GG, PK

Interpretation of the data: EA, ECC, YT, SS, JAC, GG, PK

Draft of the manuscript: EA, PK

Revision of the manuscript: EA, ECC, EY, YT, SS, JAC, GG, PK

## Acknowledgements

The authors are thankful to staff of Nefas Mewcha Hospital for their enthusiastic collaboration.

---

**Table S1:**
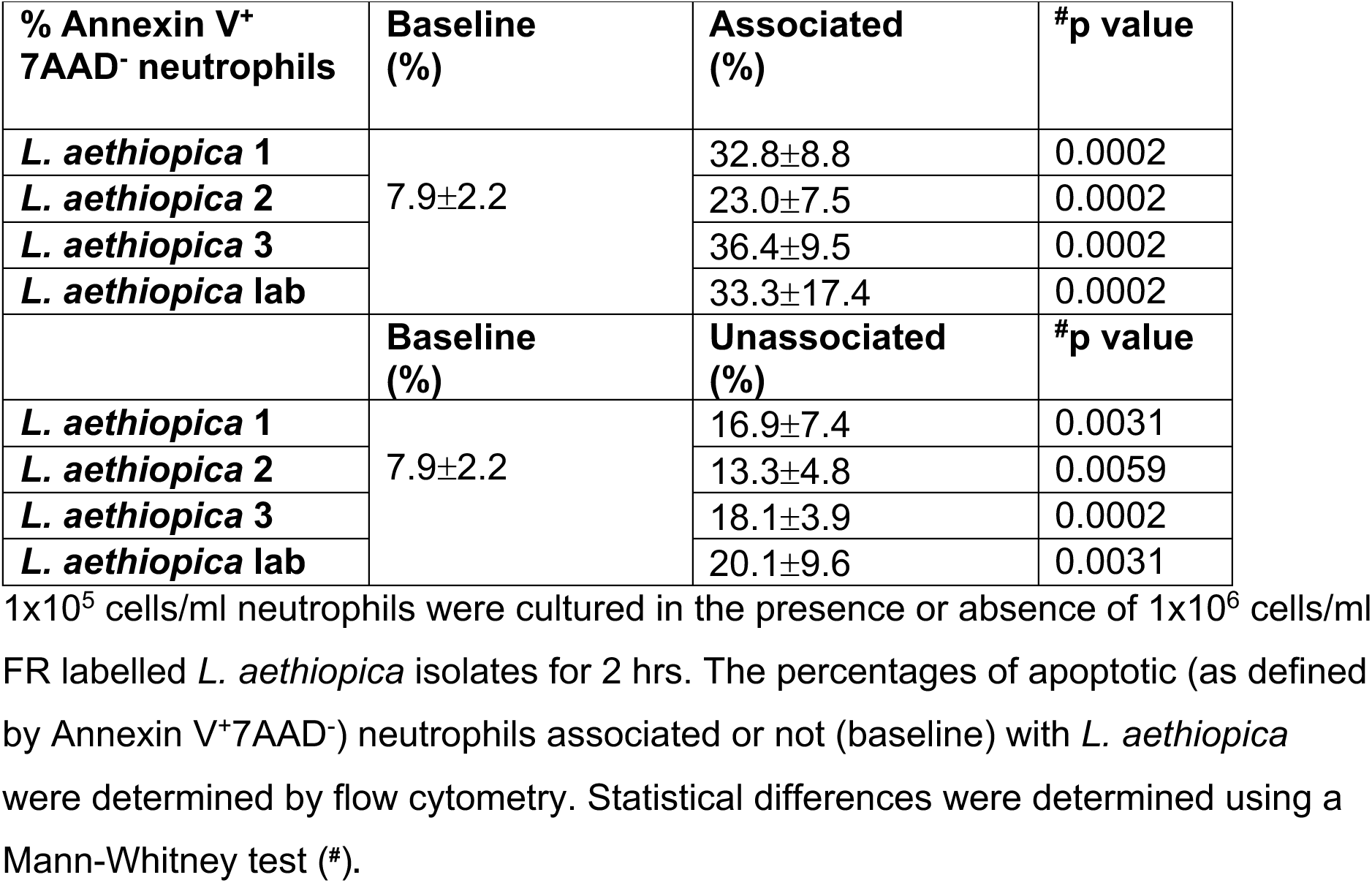
% of apoptotic neutrophils following 2 hrs of incubation.

| % Annexin V<sup>+</sup><br>7AAD<sup>-</sup> neutrophils | Baseline<br>(%) | Associated<br>(%) | #p value |
| --- | --- | --- | --- |
| *L. aethiopica* 1 | 7.9±2.2 | 32.8±8.8 | 0.0002 |
| *L. aethiopica* 2 |  | 23.0±7.5 | 0.0002 |
| *L. aethiopica* 3 |  | 36.4±9.5 | 0.0002 |
| *L. aethiopica* lab |  | 33.3±17.4 | 0.0002 |
|  | Baseline<br>(%) | Unassociated<br>(%) | #p value |
| *L. aethiopica* 1 | 7.9±2.2 | 16.9±7.4 | 0.0031 |
| *L. aethiopica* 2 |  | 13.3±4.8 | 0.0059 |
| *L. aethiopica* 3 |  | 18.1±3.9 | 0.0002 |
| *L. aethiopica* lab |  | 20.1±9.6 | 0.0031 |

1x10<sup>5</sup> cells/ml neutrophils were cultured in the presence or absence of 1x10<sup>6</sup> cells/ml FR labelled *L. aethiopica* isolates for 2 hrs. The percentages of apoptotic (as defined by Annexin V<sup>+</sup>7AAD<sup>-</sup>) neutrophils associated or not (baseline) with *L. aethiopica* were determined by flow cytometry. Statistical differences were determined using a Mann-Whitney test (#).

**Table S2:** Comparison of the % of apoptotic neutrophils between unassociated and associated neutrophils following 2hrs of co-incubation with *L. aethiopica*.

| Apoptosis | Unassociated<br>(%) | Associated<br>(%) | #p value |
| --- | --- | --- | --- |
| *L. aethiopica* 1 | 16.9±7.4 | 32.8±8.8 | 0.0022 |
| *L. aethiopica* 2 | 13.3±4.8 | 23.0±7.5 | 0.0040 |
| *L. aethiopica* 3 | 18.1±3.9 | 36.4±9.5 | 0.0003 |
| *L. aethiopica* lab | 20.1±9.6 | 33.3±17.4 | 0.1605 |

1x10<sup>5</sup> cells/ml neutrophils were co-cultured with 1x10<sup>6</sup> cells/ml FR labelled *L. aethiopica* isolates for 2 hrs. The percentages of apoptotic (as defined by Annexin V<sup>+</sup>7AAD<sup>-</sup>) neutrophils were determined by flow cytometry. Statistical differences were determined using a Mann-Whitney test (#).

**Table S3:** % of apoptotic neutrophils following 18 hrs of incubation.

| **Apoptosis** | **Baseline**<br>**(% Annexin V<sup>+</sup>**<br>**7AAD<sup>-</sup> neutrophils)** | **CD15<sup>int</sup>parasites<sup>-</sup>**<br>**(% Annexin V<sup>+</sup>**<br>**7AAD<sup>-</sup> neutrophils)** | **#p value** |
| --- | --- | --- | --- |
| *L. aethiopica* 1 | 76±6 | 76±9 | 0.9369 |
| *L. aethiopica* 2 |  | 74±9 | 0.8974 |
| *L. aethiopica* 3 |  | 80±3 | 0.1503 |
| *L. aethiopica* lab |  | 80±3 | 0.0872 |
|  | **Baseline**<br>**(% Annexin V<sup>+</sup>**<br>**7AAD<sup>-</sup> neutrophils)** | **CD15<sup>high</sup>parasites<sup>+</sup>**<br>**(% Annexin V<sup>+</sup>**<br>**7AAD<sup>-</sup> neutrophils)** | **#p value** |
| *L. aethiopica* 1 | 76±6 | 57±10 | 0.0017 |
| *L. aethiopica* 2 |  | 54±12 | 0.0006 |
| *L. aethiopica* 3 |  | 61±11 | 0.0020 |
| *L. aethiopica* lab |  | 43±12 | 0.0002 |
|  | **Baseline**<br>**(% Annexin V<sup>+</sup>**<br>**7AAD<sup>-</sup> neutrophils)** | **CD15<sup>low</sup>parasites<sup>+</sup>**<br>**(% Annexin V<sup>+</sup>**<br>**7AAD<sup>-</sup> neutrophils)** | **#p value** |
| *L. aethiopica* 1 | 76±6 | 29±16 | 0.0002 |
| *L. aethiopica* 2 |  | 32±19 | 0.0003 |
| *L. aethiopica* 3 |  | 40±14 | 0.0002 |
| *L. aethiopica* lab |  | 26±13 | 0.0002 |

1x10<sup>5</sup> cells/ml neutrophils were cultured in the presence or in the absence of 1x10<sup>6</sup> cells/ml FR labelled *L. aethiopica* isolates for 18 hrs. The percentages of apoptotic (as defined by Annexin V<sup>+</sup>7AAD<sup>-</sup>) neutrophils associated or not (baseline) with *L. aethiopica* were determined by flow cytometry. Statistical differences were determined using a Mann-Whitney test (#).

**Table S4:** Comparison of the % of apoptotic CD15^int^parasite^-^, CD15^high^parasite^+^ and CD15^low^parasite^+^ for each parasites following 18 hrs of incubation.

| ***L. aethiopica* 1** | **% apoptosis** | **^p value** |  | **#p value** |
| --- | --- | --- | --- | --- |
| CD15<sup>int</sup>parasites<sup>-</sup> | 76±9 | 0.0002 | CD15<sup>int</sup>parasites<sup>-</sup><br>vs CD15<sup>high</sup>parasites<sup>+</sup> | 0.1107 |
| CD15<sup>high</sup>parasites<sup>+</sup> | 57±10 |  | CD15<sup>int</sup>parasites<sup>-</sup><br>vs CD15<sup>low</sup>parasites<sup>+</sup> | 0.0001 |
| CD15<sup>low</sup>parasites<sup>+</sup> | 29±16 |  | CD15<sup>high</sup>parasites<sup>+</sup><br>vs CD15<sup>low</sup>parasites<sup>+</sup> | 0.1259 |
| ***L. aethiopica* 2** | **% apoptosis** | **^p value** |  | **#p value** |
| CD15<sup>int</sup>parasites<sup>-</sup> | 74±9 | 0.0009 | CD15<sup>int</sup>parasites<sup>-</sup><br>vs CD15<sup>high</sup>parasites<sup>+</sup> | 0.1367 |
| CD15<sup>high</sup>parasites<sup>+</sup> | 54±12 |  | CD15<sup>int</sup>parasites<sup>-</sup><br>vs CD15<sup>low</sup>parasites<sup>+</sup> | 0.0006 |
| CD15<sup>low</sup>parasites<sup>+</sup> | 32±19 |  | CD15<sup>high</sup>parasites<sup>+</sup><br>vs CD15<sup>low</sup>parasites<sup>+</sup> | 0.2488 |
| ***L. aethiopica* 3** | **% apoptosis** | **^p value** |  | **#p value** |
| CD15<sup>int</sup>parasites<sup>-</sup> | 80±3 | <0.0001 | CD15<sup>int</sup>parasites<sup>-</sup><br>vs CD15<sup>high</sup>parasites<sup>+</sup> | 0.0360 |
| CD15<sup>high</sup>parasites<sup>+</sup> | 61±11 |  | CD15<sup>int</sup>parasites<sup>-</sup><br>vs CD15<sup>low</sup>parasites<sup>+</sup> | <0.0001 |
| CD15<sup>low</sup>parasites<sup>+</sup> | 40±14 |  | CD15<sup>high</sup>parasites<sup>+</sup><br>vs CD15<sup>low</sup>parasites<sup>+</sup> | 0.2307 |
| ***L. aethiopica* lab** | **% apoptosis** | **^p value** |  | **#p value** |
| CD15<sup>int</sup>parasites<sup>-</sup> | 80±3 | 0.0001 | CD15<sup>int</sup>parasites<sup>-</sup><br>vs CD15<sup>high</sup>parasites<sup>+</sup> | 0.0252 |
| CD15<sup>high</sup>parasites<sup>+</sup> | 43±12 |  | CD15<sup>int</sup>parasites<sup>-</sup><br>vs CD15<sup>low</sup>parasites<sup>+</sup> | <0.0001 |
| CD15<sup>low</sup>parasites<sup>+</sup> | 26±13 |  | CD15<sup>high</sup>parasites<sup>+</sup><br>vs CD15<sup>low</sup>parasites<sup>+</sup> | 0.3846 |

1x10<sup>5</sup> cells/ml neutrophils were co-cultured with 1x10<sup>6</sup> cells/ml FR labelled *L. aethiopica* isolates for 18 hrs. The percentages of apoptotic (as defined by Annexin V<sup>+</sup>7AAD<sup>-</sup>) CD15<sup>int</sup>parasite<sup>-</sup>, CD15<sup>high</sup>parasite<sup>+</sup> and CD15<sup>low</sup>parasite<sup>+</sup> were determined by flow cytometry. Statistical differences were determined using Kruskal-Wallis (\*) and Dunn's multiple comparison (^) tests.

**Table S5:**
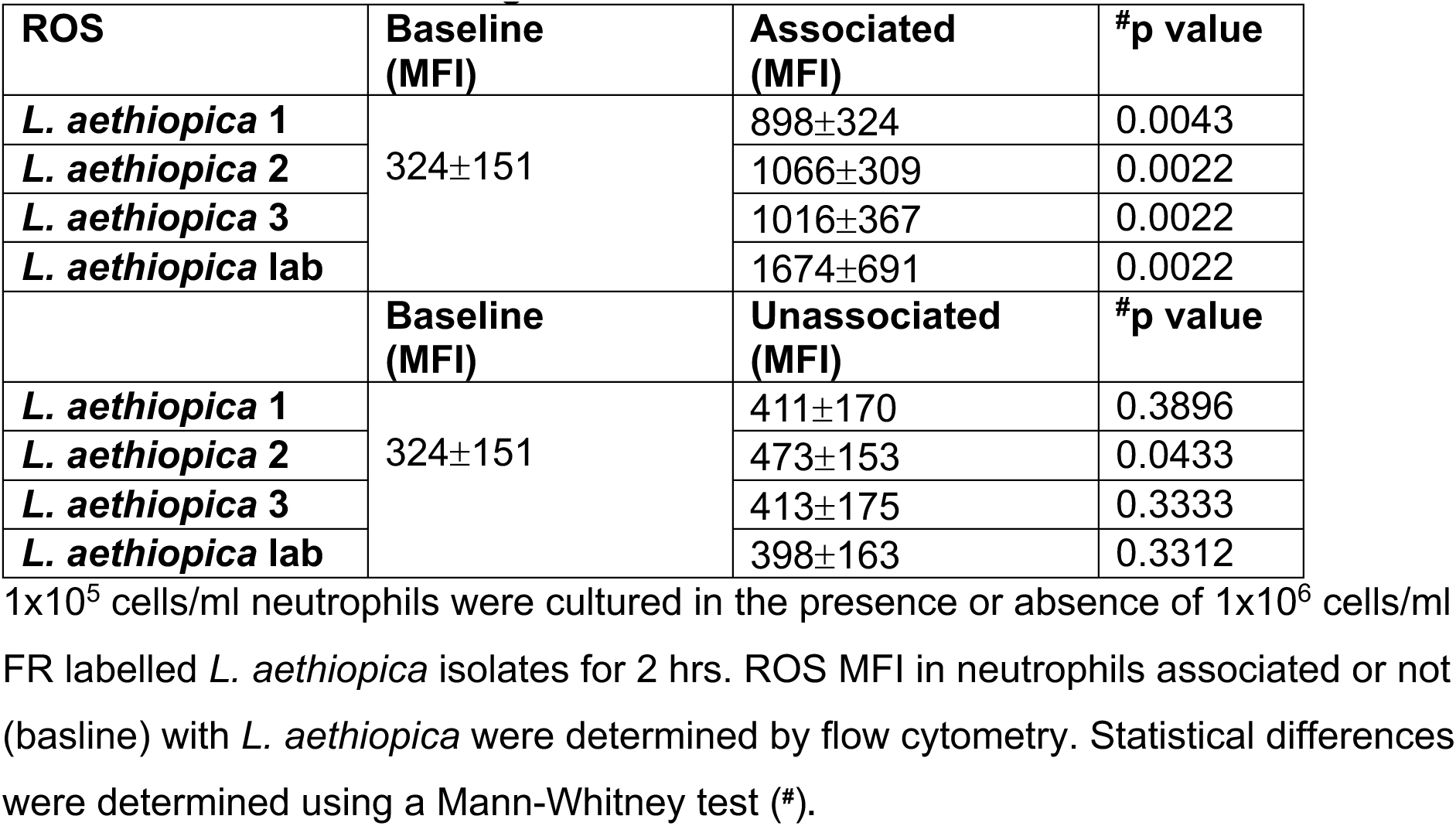
ROS MFI following 2 hrs of incubation.

**Table S6:** Comparison of ROS MFI between unassocaited and associated neutrophils following 2hrs of co-incubation with *L. aethiopica*.

| ROS | Unassociated (MFI) | Associated (MFI) | #p value |
| --- | --- | --- | --- |
| *L. aethiopica* 1 | 411±170 | 898±324 | 0.0173 |
| *L. aethiopica* 2 | 473±153 | 1066±309 | 0.0043 |
| *L. aethiopica* 3 | 413±175 | 1016±367 | 0.0043 |
| *L. aethiopica* lab | 398±163 | 1674±691 | 0.0022 |

1x10<sup>5</sup> cells/ml neutrophils were co-cultured with 1x10<sup>6</sup> cells/ml FR labelled *L. aethiopica* isolates for 2 hrs. ROS MFI were determined by flow cytometry. Statistical differences were determined using a Mann-Whitney test (#).

**Table S7:** ROS MFI in neutrophils following 18 hrs of incubation.

| **ROS** | **Baseline (MFI)** | **CD15<sup>int</sup>parasites<sup>-</sup> (MFI)** | **#p value** |
| --- | --- | --- | --- |
| *L. aethiopica* 1 | 83.33±26.88 | 80.50±25.72 | 0.9740 |
| *L. aethiopica* 2 |  | 92.83±41.56 | 0.7294 |
| *L. aethiopica* 3 |  | 98.00±37.34 | 0.4848 |
| *L. aethiopica* lab |  | 117.3±43.63 | 0.1580 |
|  | **Baseline (MFI)** | **CD15<sup>high</sup>parasites<sup>+</sup> (MFI)** | **#p value** |
| *L. aethiopica* 1 | 83.33±26.88 | 126.0±45.48 | 0.1017 |
| *L. aethiopica* 2 |  | 136.7±81.28 | 0.3095 |
| *L. aethiopica* 3 |  | 159.3±90.29 | 0.0931 |
| *L. aethiopica* lab |  | 183.2±83.60 | 0.0649 |
|  | **Baseline (MFI)** | **CD15<sup>low</sup>parasites<sup>+</sup> (MFI)** | **#p value** |
| *L. aethiopica* 1 | 83.33±26.88 | 40.00±12.43 | 0.0087 |
| *L. aethiopica* 2 |  | 30.83±8.448 | 0.0022 |
| *L. aethiopica* 3 |  | 56.17±16.18 | 0.0455 |
| *L. aethiopica* lab |  | 65.33±17.05 | 0.3701 |

1x10<sup>5</sup> cells/ml neutrophils were cultured in the presence or in the absence of 1x10<sup>6</sup> cells/ml FR labelled *L. aethiopica* isolates for 18 hrs. ROS MFI in neutrophils associated or not (baseline) with *L. aethiopica* were determined by flow cytometry. Statistical differences were determined using a Mann-Whitney test (#).

**Table S8:**
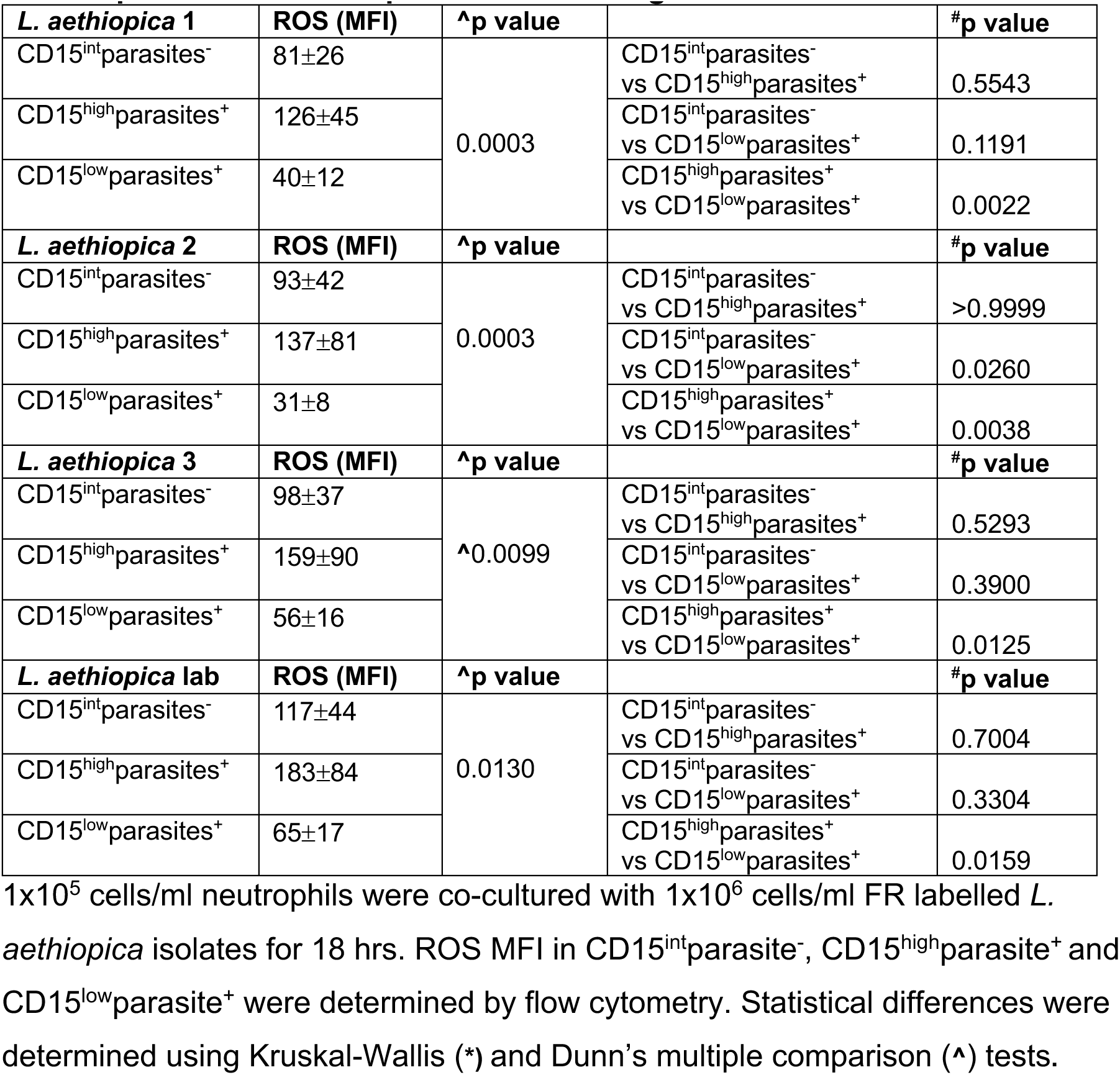
Comparison of ROS MFI in CD15^int^parasite^-^, CD15^high^parasite^+^ and CD15^low^parasite^+^ for each parasites following 18 hrs of incubation.

| ***L. aethiopica* 1** | **ROS (MFI)** | **^p value** |  | **#p value** |
| --- | --- | --- | --- | --- |
| CD15<sup>int</sup>parasites<sup>-</sup> | 81±26 | 0.0003 | CD15<sup>int</sup>parasites<sup>-</sup><br>vs CD15<sup>high</sup>parasites<sup>+</sup> | 0.5543 |
| CD15<sup>high</sup>parasites<sup>+</sup> | 126±45 |  | CD15<sup>int</sup>parasites<sup>-</sup><br>vs CD15<sup>low</sup>parasites<sup>+</sup> | 0.1191 |
| CD15<sup>low</sup>parasites<sup>+</sup> | 40±12 |  | CD15<sup>high</sup>parasites<sup>+</sup><br>vs CD15<sup>low</sup>parasites<sup>+</sup> | 0.0022 |
| ***L. aethiopica* 2** | **ROS (MFI)** | **^p value** |  | **#p value** |
| CD15<sup>int</sup>parasites<sup>-</sup> | 93±42 | 0.0003 | CD15<sup>int</sup>parasites<sup>-</sup><br>vs CD15<sup>high</sup>parasites<sup>+</sup> | >0.9999 |
| CD15<sup>high</sup>parasites<sup>+</sup> | 137±81 |  | CD15<sup>int</sup>parasites<sup>-</sup><br>vs CD15<sup>low</sup>parasites<sup>+</sup> | 0.0260 |
| CD15<sup>low</sup>parasites<sup>+</sup> | 31±8 |  | CD15<sup>high</sup>parasites<sup>+</sup><br>vs CD15<sup>low</sup>parasites<sup>+</sup> | 0.0038 |
| ***L. aethiopica* 3** | **ROS (MFI)** | **^p value** |  | **#p value** |
| CD15<sup>int</sup>parasites<sup>-</sup> | 98±37 | ^0.0099 | CD15<sup>int</sup>parasites<sup>-</sup><br>vs CD15<sup>high</sup>parasites<sup>+</sup> | 0.5293 |
| CD15<sup>high</sup>parasites<sup>+</sup> | 159±90 |  | CD15<sup>int</sup>parasites<sup>-</sup><br>vs CD15<sup>low</sup>parasites<sup>+</sup> | 0.3900 |
| CD15<sup>low</sup>parasites<sup>+</sup> | 56±16 |  | CD15<sup>high</sup>parasites<sup>+</sup><br>vs CD15<sup>low</sup>parasites<sup>+</sup> | 0.0125 |
| ***L. aethiopica* lab** | **ROS (MFI)** | **^p value** |  | **#p value** |
| CD15<sup>int</sup>parasites<sup>-</sup> | 117±44 | 0.0130 | CD15<sup>int</sup>parasites<sup>-</sup><br>vs CD15<sup>high</sup>parasites<sup>+</sup> | 0.7004 |
| CD15<sup>high</sup>parasites<sup>+</sup> | 183±84 |  | CD15<sup>int</sup>parasites<sup>-</sup><br>vs CD15<sup>low</sup>parasites<sup>+</sup> | 0.3304 |
| CD15<sup>low</sup>parasites<sup>+</sup> | 65±17 |  | CD15<sup>high</sup>parasites<sup>+</sup><br>vs CD15<sup>low</sup>parasites<sup>+</sup> | 0.0159 |

1x10<sup>5</sup> cells/ml neutrophils were co-cultured with 1x10<sup>6</sup> cells/ml FR labelled *L. aethiopica* isolates for 18 hrs. ROS MFI in CD15<sup>int</sup>parasite<sup>-</sup>, CD15<sup>high</sup>parasite<sup>+</sup> and CD15<sup>low</sup>parasite<sup>+</sup> were determined by flow cytometry. Statistical differences were determined using Kruskal-Wallis (\*) and Dunn's multiple comparison (^) tests.

## S FIGURE LEGENDS

**FIGURE S1:**
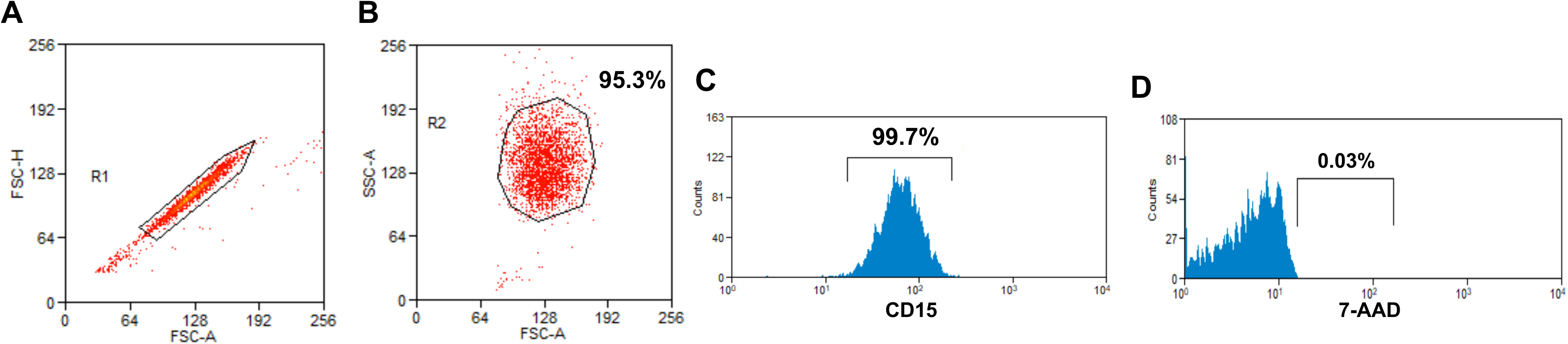
Gating strategy for neutrophils after purification from peripheral blood mononuclear cells. Following neutrophils purification, doublets were excluded (**A**), neutrophils were identified by FSC and SSC (**B**) and anti-CD15 mAb (**C**) the number of dead cells as defined by 7-AAD positive cells were determined by flow cytometry (**D**).

**FIGURE S2:**
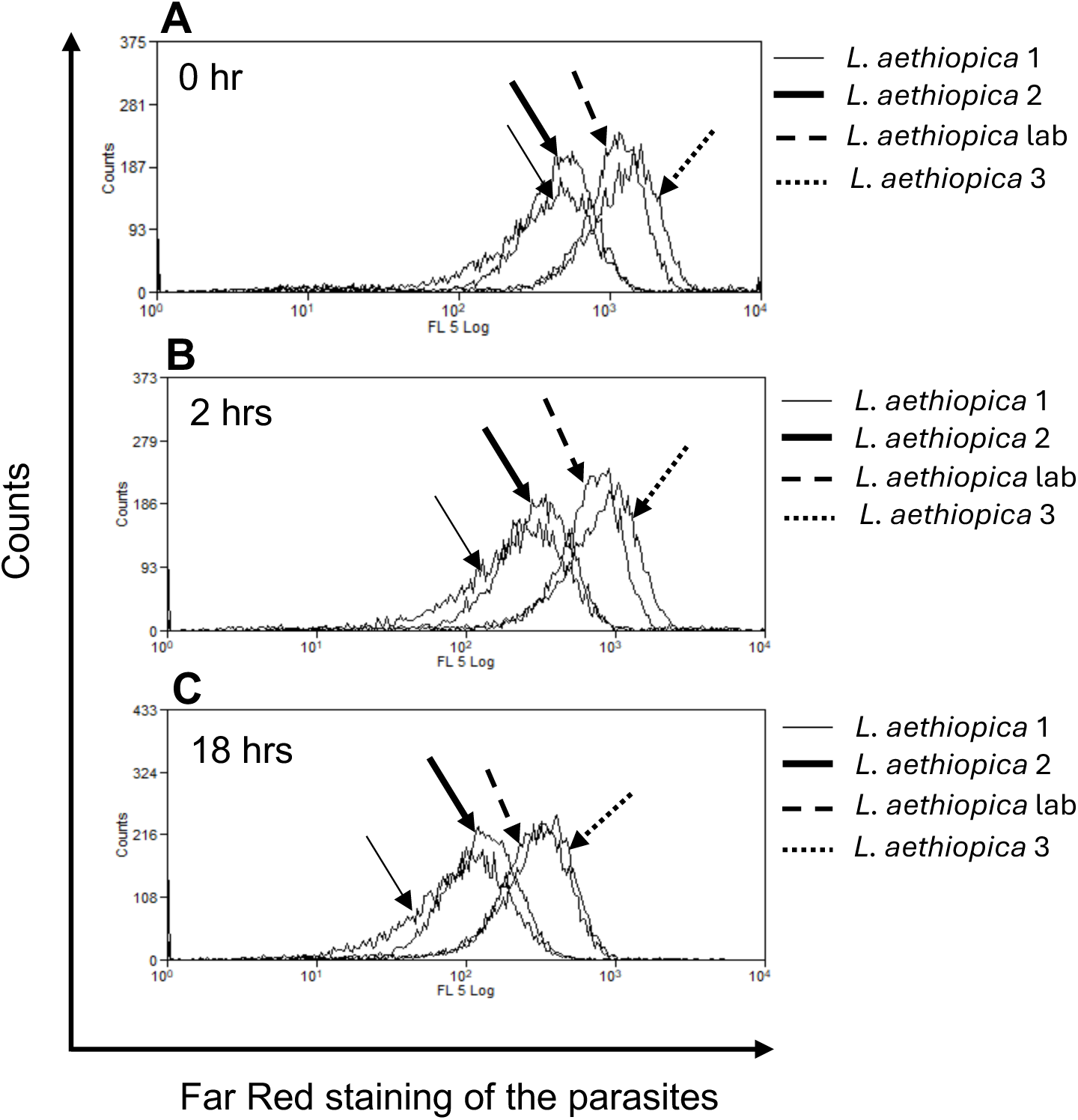
Labelling of *L. aethiopica* with Far Red. The different isolates of *L. aethiopica* were labelled with Far Red (FR) as described in Materials and Methods and the intensity of FR for each isolate was measured by flow cytometry directly after labelling (**A**), after 2hrs (**B**) and 18hrs (**C**) at 37°C.

**FIGURE S3:**
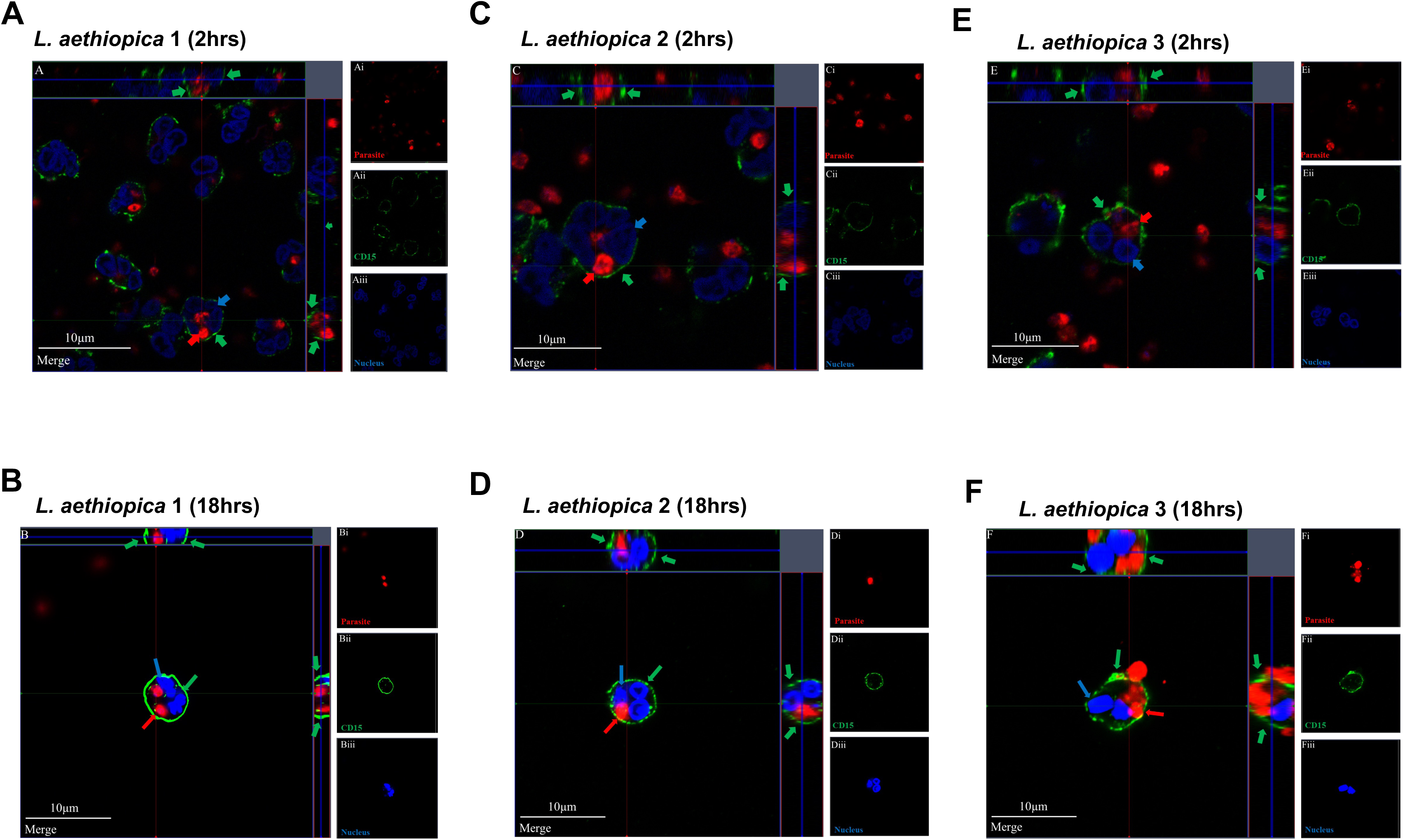
Internalisation of *L. aethiopica* by neutrophils. 1×10^5^ cells/ml neutrophils were co-cultured with 1×10^6^ cells/ml FR labelled parasites. Cells were stained as described in Materials and Methods. Confocal images were taken for *L. aethiopica* 1 after 2 hrs (**A**) and 18 hrs (**B**), *L. aethiopica* 2 after 2 hrs (**C**) and 18 hrs (**D**) and *L. aethiopica* 3 after 2 hrs (**E**) and 18 hrs (**F**). The red arrows point to the parasite (FR), the green arrows to the CD15 (Alexa Fluor 555) and the blue arrow to the nucleus (DAPI). These are representative images of at least three independent experiments. One representative image of at least three independent experiments is shown.

**FIGURE S4:**
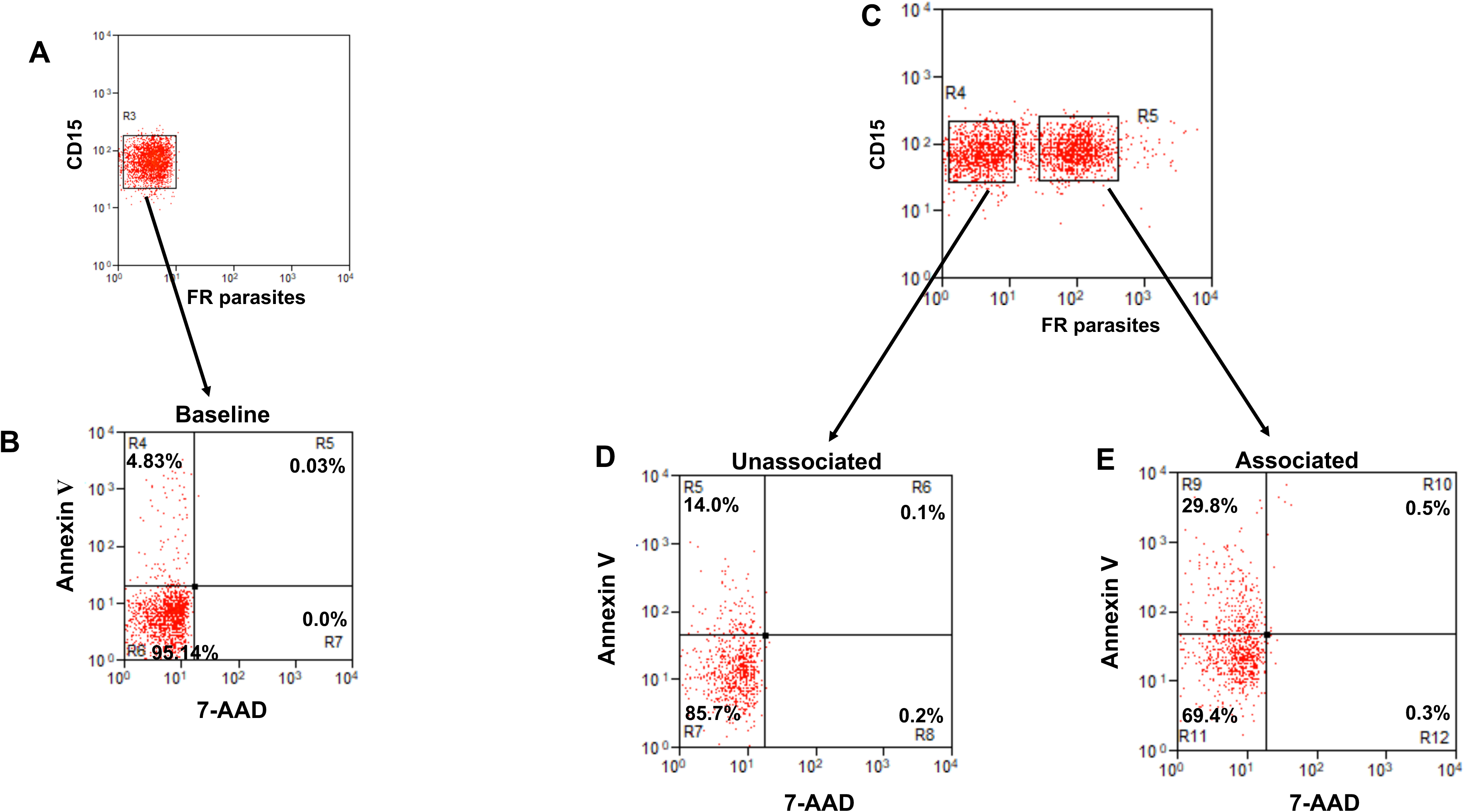
Percentages of apoptotic neutrophils after 2 hours: gating strategy. 1×10^5^ cells/ml neutrophils were cultured alone (**A** and **B**) or with 1×10^6^ cells/ml FR labelled *L. aethiopica* isolates for 2 hrs (**C, D and E**). **A**. Dot plot of CD15^+^ cells (Gate R3). **B**. Dot plot showing the percentages of Annexin V^+^ and 7-AAD^+^ in CD15^+^ cells. (**C**) Dot plot showing neutrophils unassociated (gate R4) and associated (gate R5) with *L. aethiopica*. (**D**) Dot plot showing the percentages of Annexin V^+^ and 7-AAD^+^ in unassociated CD15^+^ cells. (**E**) Dot plot showing the percentages of Annexin V^+^ and 7-AAD^+^ in associated CD15^+^ cells.

**FIGURE S5:**
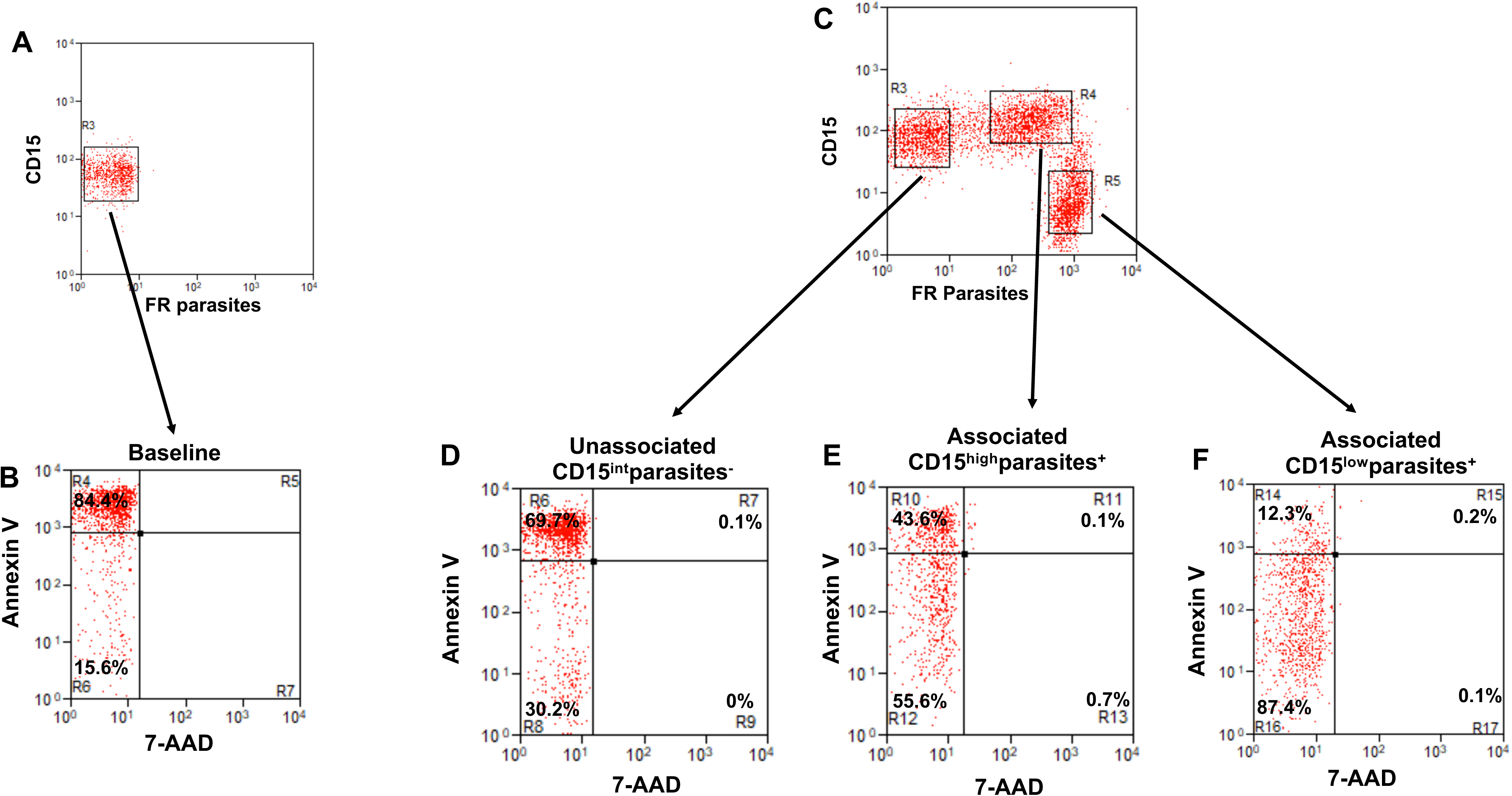
Percentages of apoptotic neutrophils after 18 hours: gating strategy. 1×10^5^ cells/ml neutrophils were cultured alone (**A** and **B**) or with 1×10^6^ cells/ml FR labelled *L. aethiopica* isolates for 18 hrs (**C, D, E and F**). **A**. Dot plot of CD15^+^ cells (Gate R3). **B**. Dot plot showing the percentages of Annexin V^+^ and 7-AAD^+^ in CD15^+^ cells. (**C**) Dot plot showing neutrophils unassociated: CD15^int^parasite^-^ (gate R3) and associated: CD15^high^parasite^+^ (gate R4) and CD15^low^parasite^+^ (gate R5) with *L. aethiopica*. (**D**) Dot plot showing the percentages of Annexin V^+^ and 7-AAD^+^ in CD15^int^parasite^-^. (**E**) Dot plot showing the percentages of Annexin V^+^ and 7-AAD^+^ in CD15^high^parasite^+^. (**F**) Dot plot showing the percentages of Annexin V^+^ and 7-AAD^+^ in CD15^low^parasite^+^.

**FIGURE S6:**
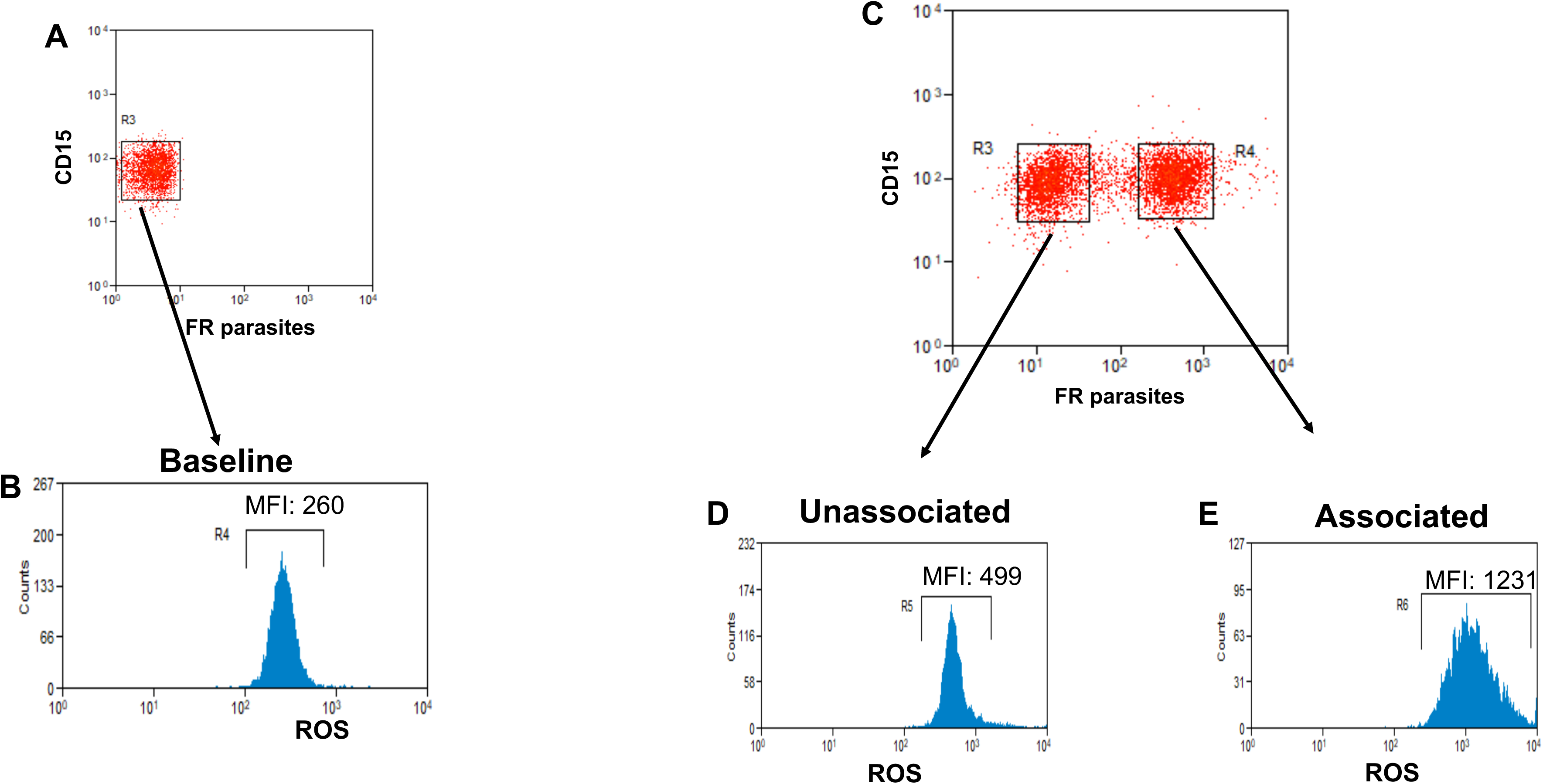
ROS MFI in neutrophils after 2 hours: gating strategy. 1×10^5^ cells/ml neutrophils were cultured alone (**A** and **B**) or with 1×10^6^ cells/ml FR labelled *L. aethiopica* isolates for 2 hrs (**C, D and E**). **A**. Dot plot of CD15^+^ cells (Gate R3). **B**. Histogram showing the ROS MFI in CD15^+^ cells. (**C**) Dot plot showing neutrophils unassociated (gate R3) and associated (gate R4) with *L. aethiopica*. (**D**) Histogram showing the ROS MFI in unassociated CD15^+^ cells. (**E**) Dot plot showing Histogram showing the ROS MFI in associated CD15^+^ cells.

**FIGURE S7:**
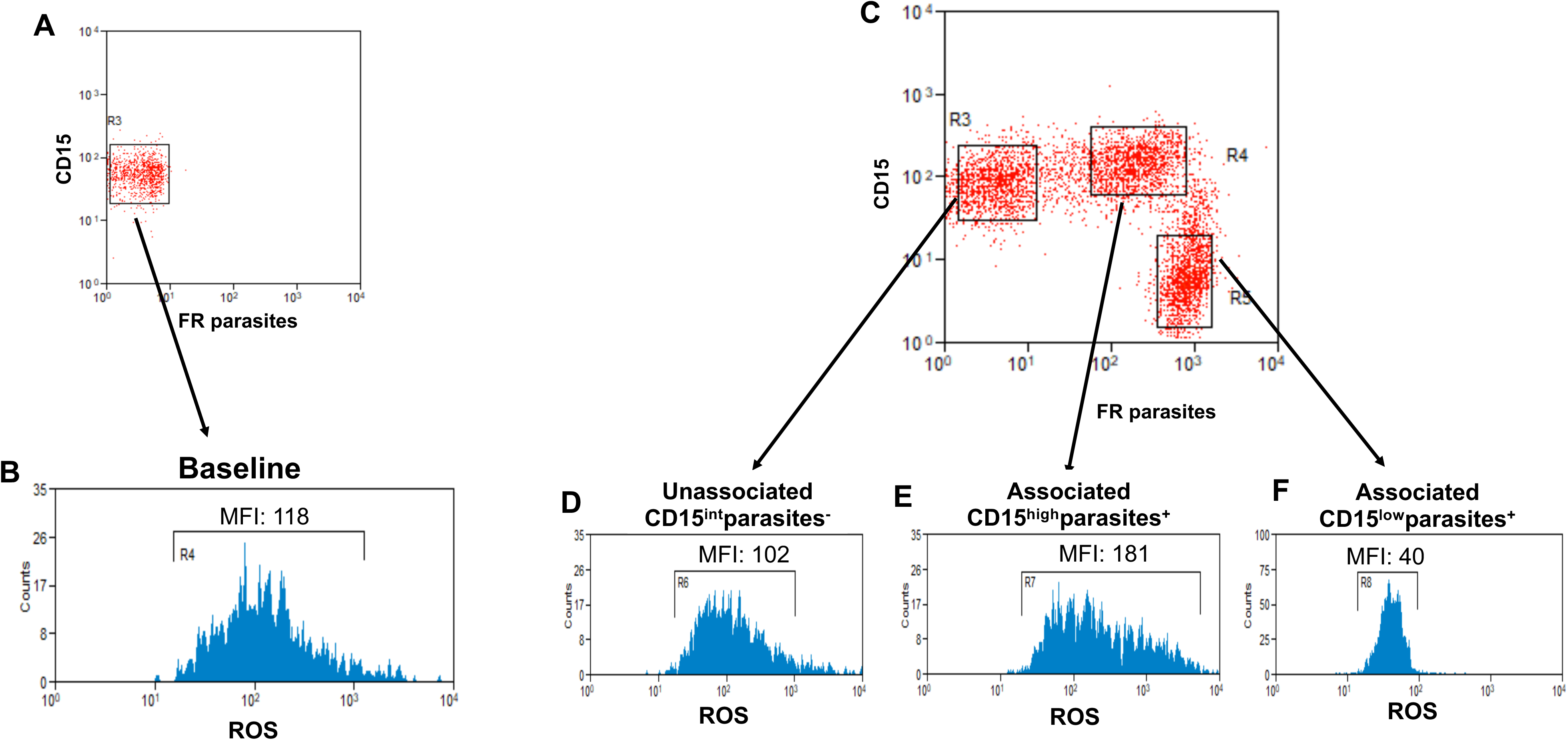
ROS MFI in neutrophils after 18 hours: gating strategy. 1×10^5^ cells/ml neutrophils were cultured alone (**A** and **B**) or with 1×10^6^ cells/ml FR labelled *L. aethiopica* isolates for 18 hrs (**C, D, E and F**). **A**. Dot plot of CD15^+^ cells (Gate R3). **B**. Histogram showing the ROS MF in CD15^+^ cells. (**C**) Dot plot showing neutrophils unassociated: CD15^int^parasite^-^ (gate R3) and associated: CD15^high^parasite^+^ (gate R4) and CD15^low^parasite^+^ (gate R5) with *L. aethiopica*. (**D**) Histogram showing the ROS MFI in CD15^int^parasite^-^. (**E**) Histogram showing the ROS MFI CD15^high^parasite^+^. (**F**) Histogram showing the ROS MFI in CD15^low^parasite^+^.

